# Combined TIRF and 3D Super-Resolution Microscopy for Nanoscopic Spatiotemporal Characterization of Adhesion Molecules on Microvilli

**DOI:** 10.1101/2025.08.26.672391

**Authors:** Abdullah Alghamdi, Mansour M. Aldehaiman, Maged F. Serag, Shuho Nozue, Ioana-Andreea Ciocanaru, Jasmeen S. Merzaban, Satoshi Habuchi

## Abstract

The homing of hematopoietic stem/progenitor cells (HSPCs) and leukemic cells is a multistep process governed by complex spatiotemporal interactions between adhesion molecules under shear stress. While the molecular and biological mechanisms of this process have been extensively studied, the precise spatial and temporal organization of adhesion molecules that influences homing efficiency remains relatively poorly understood. In particular, the roles of the cell surface topography and its morphological changes during homing in shaping the spatial organization of adhesion molecules remain elusive. This is partly due to the lack of imaging techniques that simultaneously capture both nanoscopic cell surface morphology and the spatial distribution of the adhesion molecules. Here, we develop a microfluidics-based super-resolution (SR) imaging platform that enables the three-dimensional (3D) mapping of the cell surface morphology and the spatial distribution of the adhesion molecules during HSPC and leukemic cell rolling by integrating total internal reflection fluorescence microscopy (TIRFM) with single-molecule localization microscopy (SMLM). We reconstruct the cell surface morphology, which is critical to the homing, using TIRFM, and precisely overlay the spatial distribution of adhesion molecules, including CD44, PSGL-1, and actin cytoskeleton, determined by 3D-SMLM, on the topographic map. We show distinct nanoscopic localizations of adhesion molecules on the microvilli of HSPCs/leukemic cells and their reorganization under shear stress during cell rolling, at a spatial resolution of approximately 30 nm. The approach offers a powerful means to elucidate the complicated interplay between cell surface morphology and ligand-receptor interactions.

**TOC graphic:** 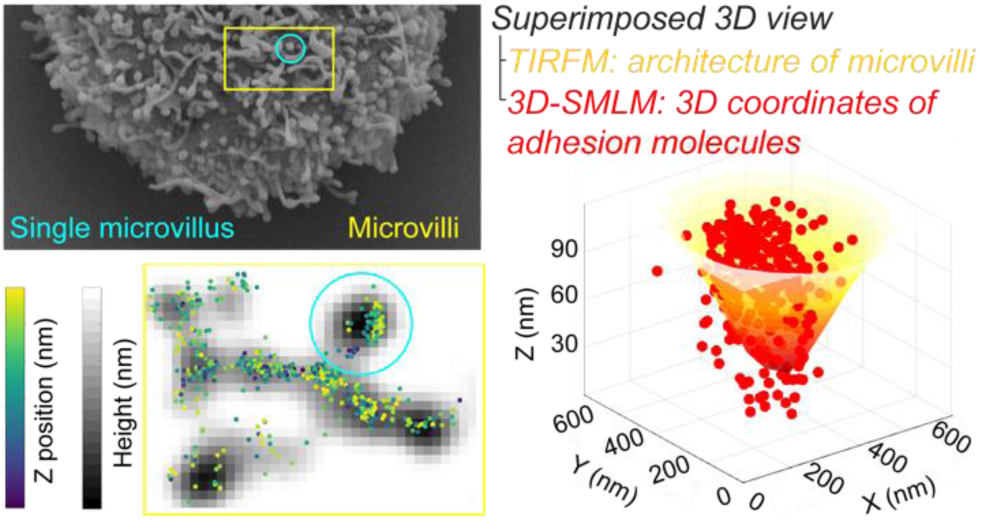

## INTRODUCTION

Homing is a fundamental biological phenomenon essential for hematopoiesis, ensuring the generation and replenishment of the blood and immune systems throughout life.^1,2^ Understanding the homing mechanisms of hematopoietic stem/progenitor cells (HSPCs) is critical for elucidating physiological processes that rely on delivering these cells to specific sites within the body.^3,4^ HSPC homing from peripheral blood to bone marrow follows a well-coordinated multistep cascade, including tethering, rolling, firm adhesion, and transmigration (Figure S1).^5^ These sequential steps are essential for efficient homing, as they rely on intricate adhesion molecule interactions that operate within the dynamic environment of blood flow and shear stress^6,7^.

Despite extensive research on the molecular aspects of cellular adhesion, the spatiotemporal dynamics of these processes and their relationship to cellular architecture remain poorly understood. A key structural feature of HSPCs is their microvilli, finger-like membranous projections supported by the actin cytoskeleton, that influence cell migration.^8^ Microvilli exhibit notable flexibility and dynamic remodeling, and studies suggest they facilitate HSPC capture on endothelial surfaces and the formation and elongation of tethers during homing.^9-15^ However, their precise physiological functions, structural biophysics, and the role of the spatial adhesion-molecules organization remain elusive. Investigating the spatial distribution of selectin ligands on HSPC microvilli is, therefore, essential for understanding the mechanisms of stable rolling and efficient homing.

Morphological studies using electron microscopy (EM) have revealed nanoscopic alterations in cell surface morphology during early stages of homing, including changes in microvilli length, width, and adhesion-related structure.^16,17^ However, traditional EM approaches, such as immunogold labeling, face limitations, including sample distortion and artifacts.^18,19^ Fluorescence-based studies often treat the HSPC membrane as a flat 2D sheet, neglecting its complex 3D microvillar architecture.^16, 20-23^ These limitations underscore the need for advanced experimental approaches capable of simultaneously capturing structural biophysics, protein dynamics, and spatiotemporal interactions in live cells under physiologically relevant conditions. Advanced fluorescence imaging has begun to unravel the nanoscale organization of selectin ligands such as PSGL-1 and CD44 on microvilli, which are crucial for adhesion, tethering, and rolling of HSPCs and leukemic cells, with E-selectin interactions being strengthened by structural features beyond its lectin domain^24^. Studies have demonstrated that selectin ligands exist as preformed nanoclusters tens of nanometers in size, even before rolling.^25-27^ SR imaging further reveals that under shear stress, CD44 nanoclusters reorganize into elongated network-like structures, greatly increasing their membrane footprint.^8^ This reorganization parallels observations in neutrophils rolling on endothelial surfaces.^8, 9, 28^ However, the observed elongation of CD44 exceeds expected microvilli dimensions, prompting a reevaluation of current models of membrane protrusions and their role in adhesion dynamics.^29-31^

These findings emphasize a central role of microvilli in early HSPC homing, particularly in adhesion molecule clustering and tether formation. Yet, the relationship between ligand reorganization and rolling-associated morphological changes remains poorly understood. Previous studies have combined total internal reflection fluorescence microscopy (TIRFM) with super-resolution (SR) imaging to generate 3D surface maps alongside 2D projections of ligand distribution on the membrane.^32-34^ However, these methods fall short in directly resolving the 3D localization of ligand molecules. Additionally, heterogeneous dye incorporation can cause nonuniform signal intensities, compromising axial z-mapping accuracy.^35^ TIRFM also demands precise calibration of illumination angles and evanescent field parameters for each imaging session; even minor misalignments can lead to significant errors in axial localization, distorting topography measurements and protein-location interpretation.^36^ Collectively, these limitations highlight the need for advanced imaging techniques capable of capturing the spatiotemporal dynamics of membrane-associated molecules in 3D.

Here, we present a novel microfluidics-based experimental platform that enables nanoscopic three-dimensional (3D) characterization of both cell morphology and adhesion molecule distribution during rolling. By integrating total internal reflection fluorescence microscopy (TIRFM)-based 3D mapping of the cell surface morphology^37,38^ with 3D single-molecule localization microscopy (3D-SMLM), we reconstruct the 3D topography of fluorescently labeled HSPCs rolling on a selectin-coated microfluidic chamber and simultaneously determine the 3D spatial distribution of key adhesion molecules (Figure 1). This approach allows precise localization of selectin ligands on individual microvilli and assessment of their reorganization under shear flow. Using this method, we reveal distinct nanoscopic localizations of adhesion molecules on HSPCs/leukemic cell microvilliand their reorganization during cell rolling.

**Figure 1.**
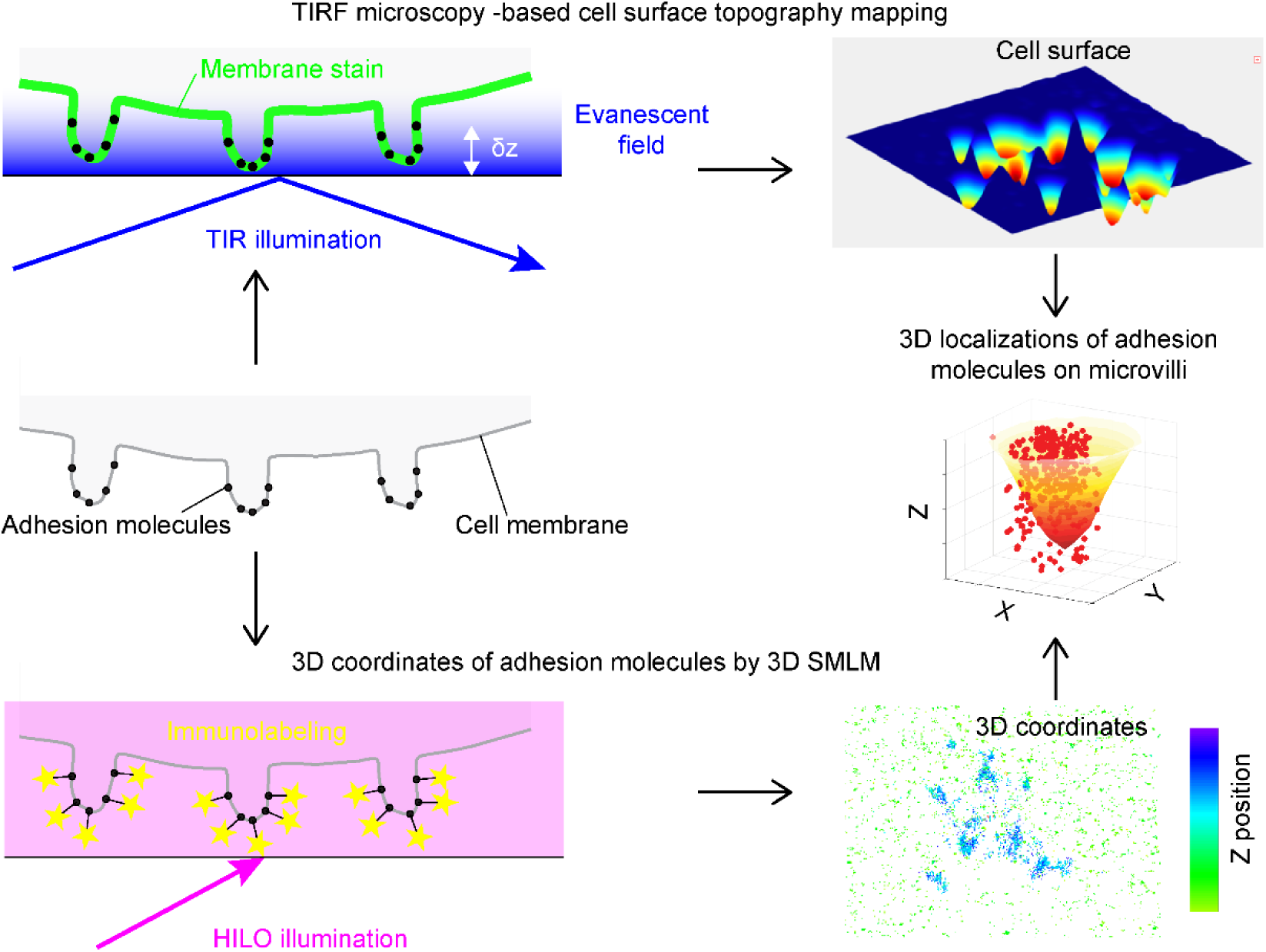
Schematic illustration of the combined TIRFM and 3D-SMLM. 3D cell surface topography was obtained by the depth-dependent fluorescence intensity of the membrane stain by TIRF microscopy with TIR illumination at 640 nm, whereas 3D coordinates of the immunolabeled adhesion molecules were determined by 3D-SMLM with highly inclined and liminated optical sheet (HILO) illumination at 488 nm.

## RESULTS AND DISCUSSION

### Design of imaging experiments and image processing pipeline

To investigate the nanoscale localization of membrane-associated adhesion molecules on the microvilli of HSPCs/leukemic cells, we determined the 3D topography of the cell surface, in particular, the microvilli structure, and overlaid 3D coordinates of the adhesion molecules on the topographic map. This was done by combining TIRFM for the topographic mapping and 3D-SMLM for the determination of the 3D coordinates (Figure S2). In the previous studies on the combined TIRFM/2D-SMLM, 2D coordinates of membrane-associated molecules were overlaid with a TIRFM-based 3D topographic map of the cell surface.^32, 34, 39^ This experimental configuration does not provide exact locations and distributions of the membrane-associated molecules in the 3D space (i.e., on the microvilli). In this study, we achieved 3D localization of adhesion molecules on the microvilli by combining TIRFM and 3D-SMLM.

We integrated the combined TIRFM/3D-SMLM into a microfluidics-based cell rolling assay platform to characterize the selectin-mediated morphology change of the microvilli and the associated change in the localizations/distributions of the adhesion molecules. TIRFM/3D-SMLM imaging experiment was conducted on control cells or the cells rolled on the E-selectin surface by labeling the cell membrane using a membrane stain and immunolabeling the adhesion molecules. Unlike the previous TIRFM/2D-SMLM imaging studies, we captured TIRFM and 3D-SMLM images sequentially by switching the illumination mode from the TIR to Epi (or HILO) on two EM-CCD cameras, which guaranteed the acquisition of the TIRFM and 3D-SMLM images under the optimum conditions.

Overlaying the two 3D data (i.e., 3D topographic map of the cell surface and 3D coordinates of the adhesion molecules) enabled us to clearly distinguish between vertically oriented microvilli (whose 3D structure can be accurately reconstructed using the TIRFM-based topography mapping) and horizontally oriented microvilli (whose 3D structure cannot be accurately reconstructed using the TIRF-based topography mapping, which could cause a misinterpretation of the localization/distribution of the adhesion molecules). The structural similarity (SSIM) index analysis enabled us to evaluate the spatial localization of the adhesion molecules on the microvilli in 3D space. The robust data acquisition and image-processing pipeline we developed in this study (Figure 2) allowed reliable analysis of the spatial organization of adhesion molecules on microvilli (see below). The integration of the combined TIRFM/3D-SMLM with the microfluidics-based cell rolling assay platform enabled the nanoscopic characterization of the spatial re-organization of the microvilli structure and the adhesion molecules occurring during cell rolling.

**Figure 2.**
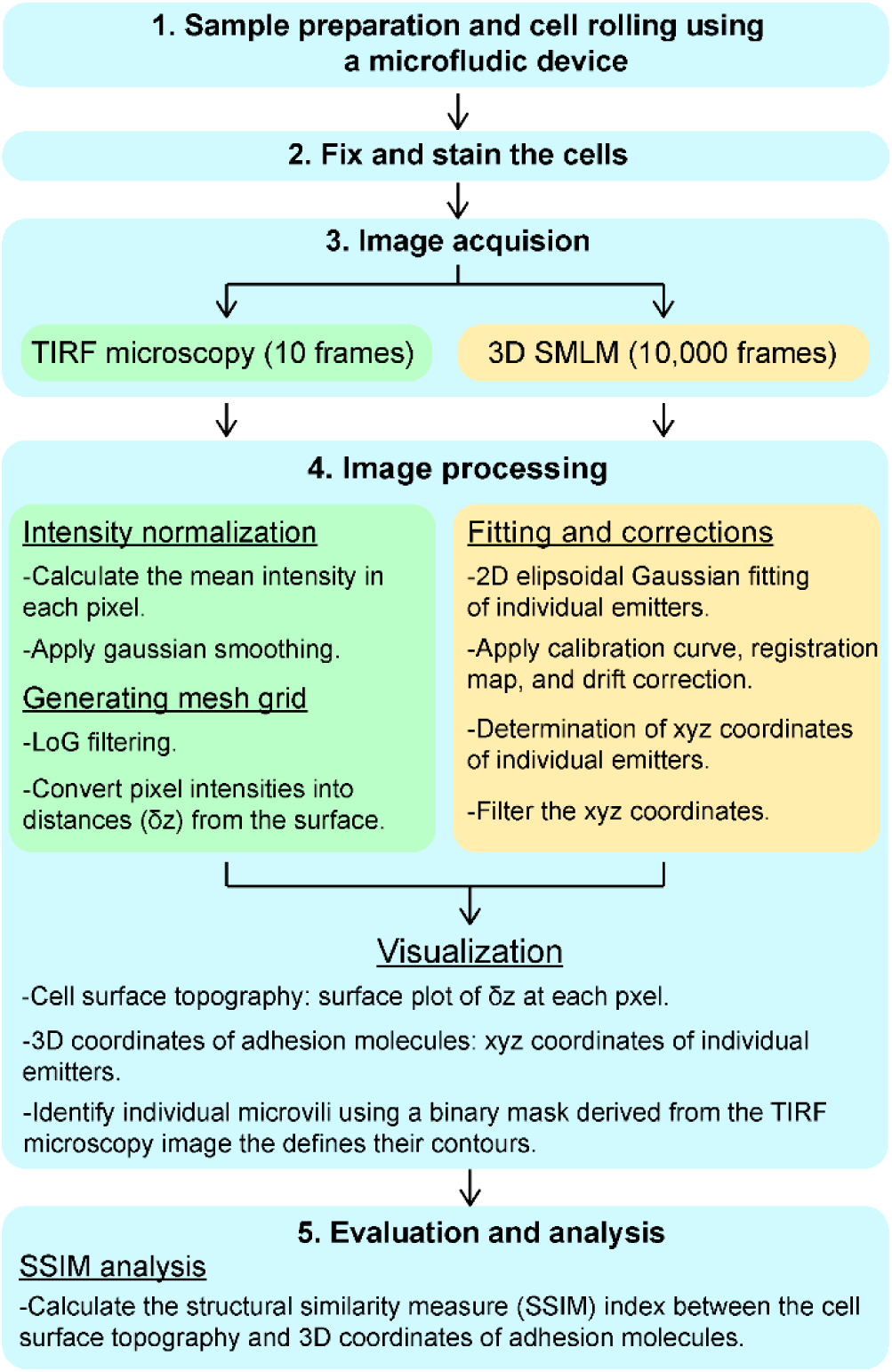
Summary of the procedures for image acquisition, processing, and analysis for combining TIRFM and 3D-SMLM datasets that provide 3D coordinates of adhesion molecules overlaid with microvilli on the cell surface.

### 3D mapping of cell surface morphology using TIRFM

3D cell surface topography was determined by depth-dependent fluorescence intensity under the TIR illumination, similar to the previous studies.^33, 34, 37^ Since the intensity of the evanescent field decays exponentially from the interface (i.e., glass-buffer interface), the nanoscopic depth profile of the cell surface is determined by fluorescence intensity variation observed for a cell stained uniformly with a membrane stain. We note that the spatial resolution of TIRFM in the XY plane is diffraction-limited.

The uniform staining of the cell membrane is essential for accurately mapping the 3D topography of microvilli. We employed two different fixable cell surface stains, DiD-Vybrant and MemBrite Fix 640 (Figures 3a, b, and S3). DiD-Vybrant is a highly lipophilic polymethine dye that is incorporated into cell membranes, and a similar dye has been used for the TIRFM-based topographic mapping of the cell surface.^40^ On the other hand, MemBrite Fix 640 reacts covalently with cell surface proteins upon accumulation in the cell membrane.^41^ We assessed the staining uniformity of these membrane stains by labeling live KG1a cells, fixing them, and mounting them in a silane-coated microfluidic chamber to preserve cell surface integrity and minimize substrate-induced distortions. The cross-sectional fluorescence images of the membrane-stained KG1a cells showed that MemBrite Fix 640 provided uniform labeling (Figures 3a and S3a, b), whereas DiD-Vybrant exhibited non-uniform staining with dye aggregates disrupting fluorescence homogeneity (Figures 3b and S3c, d). The fluorescence images of the bottom cell surface captured under the TIR illumination showed that the DiD-Vybrant-stained cells revealed dye aggregates away from and underneath the bottom membrane (Figure S3c), whereas MemBrite Fix 640-stained cells displayed a clear background with no dye aggregation, allowing for precise identification of microvilli structures (Figure S3a). These results demonstrate that MemBrite Fix 640 offers superior uniform membrane labeling, making it the preferred stain choice for 3D-TIRFM-based topographic imaging of HSPC microvilli.

**Figure 3.**
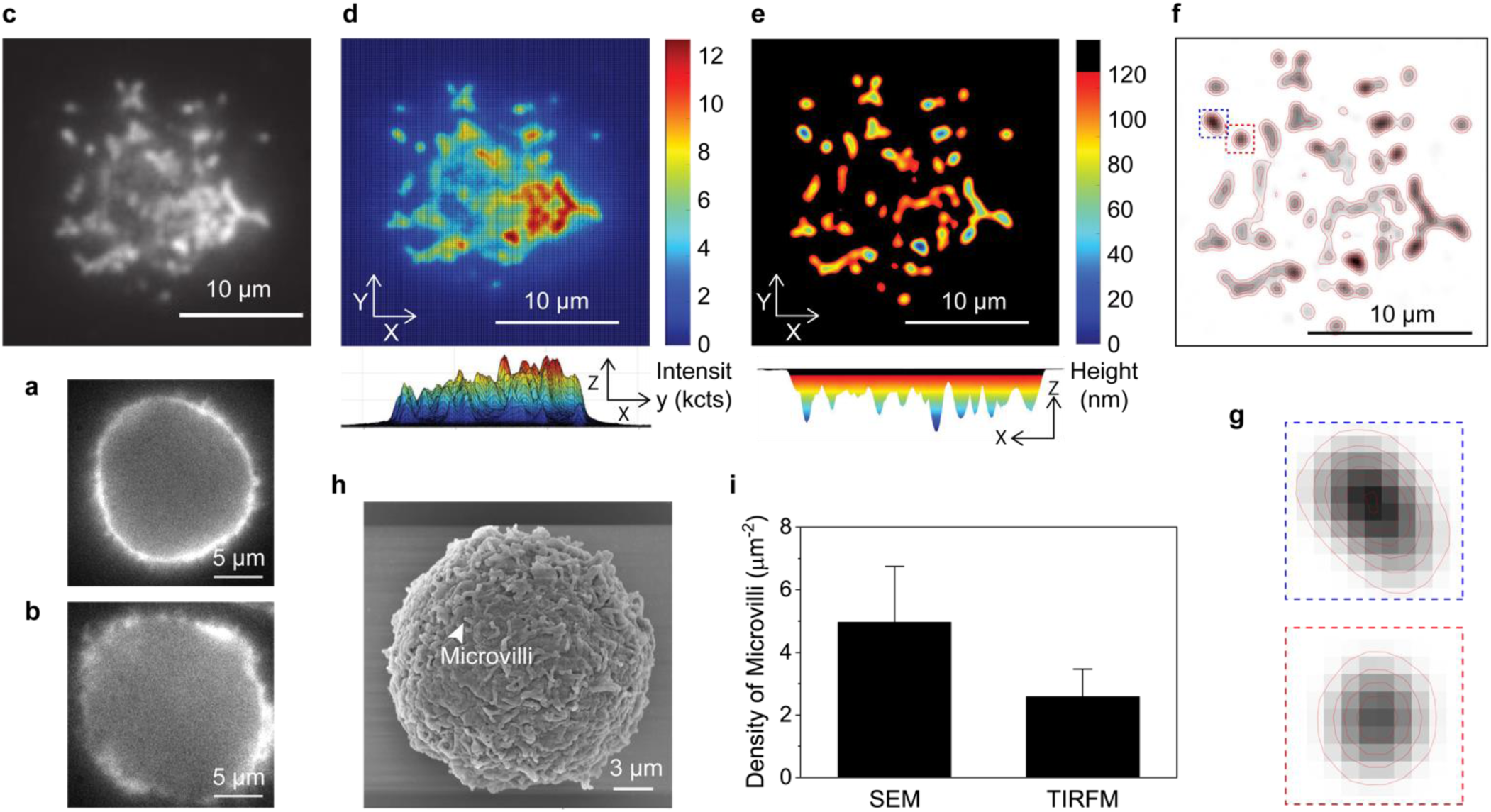
Surface topography of KG1a cells reconstructed by TIRF microscopy. (a, b) Fluorescence images of KG1a cells stained by (a) MemBrite FX640 and (b) DiD-Vybrant membrane stains, captured using wide-field epi-illumination microscopy. (c) Fluorescence image of the bottom surface of the MemBrite FX640-stained KG1a cell, captured by TIRFM. (d) Filtered image of (c) using Laplacian of Gaussian (LoG) filter; Bottom view (top panel) and side view (bottom panel). The color bar indicates fluorescence intensity values. (e) 3D topographic map reconstructed from the fluorescence image shown in (d) using equation 1; Bottom view (top panel) and side view (bottom panel). The color bar represents the distance from the glass surface. (f) Contours of individual microvilli in the topographic map shown in (e), obtained using an edge detection algorithm applied to the binary mask generated by thresholding the image. (g) Enlarged views of the regions highlighted by the rectangles in (f). δz values are red color-coded, with a step size of 10 nm. (h) SEM image of a KG1a cell, highlighting microvilli structures. (i) Comparative analysis of the average number of microvilli per µm² detected using SEM and TIRFM.

An accurate determination of the penetration depth of the evanescent field is also essential for the accurate conversion of the fluorescence intensity image into 3D cell surface topography. We experimentally verified the theoretically calculated penetration depth by the incident angle-dependent TIRF intensities of the Alexa Fluor 647 dyes in an aqueous solution (Figures S4 and S5, see Supporting Texts 1 and 2). The fluorescent intensity obtained at varied TIR illumination angles between 63 and 76 degrees fitted well with the theoretically predicted angle-dependent integrated intensity (Equation S2, Figure S5a).^42^ We observed a deviation from the theoretical prediction at the TIR angles very close to the critical angle (61 degrees, Figure S5b), probably due to either an error in the estimation of the incident angle of the excitation laser or an imperfect collimation of the excitation laser. Since our primary interest is the microvilli structure on the cell surface, we selected the TIR illumination angle of 67 degrees, corresponding to a penetration depth of 120 nm, for the TIRFM-based 3D cell surface topography mapping. This illumination condition assures that the penetration depth and, therefore, the conversion of the intensity variation into the 3D cell surface topography can be done accurately using the TIRF theory.

We captured 3D-TIRFM images of Membrite Fix 640-stained, fixed KG1a cells adhered to a clean, silane-coated glass surface with an incident angle of 67 degrees (Figure 3c). We converted the intensity images into the 3D morphology of the cell surface using a MATLAB-based image-processing pipeline (see Materials and Methods for details) in a way similar to the previous studies on the TIRFM-based cell surface topography mapping.^33, 34, 37^ Briefly, we first applied a Laplacian of Gaussian (LoG) filter (Gaussian σ = 0.5; kernel size = 10 × 10) with additional Gaussian smoothing to enhance the detection of microvilli while reducing background noise from the cell body (Figure 3d).^33^ We then converted the fluorescence intensities (*I*_z_) into relative distance values (δz) from the glass substrate using the following equation,

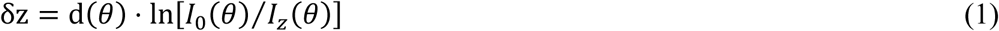

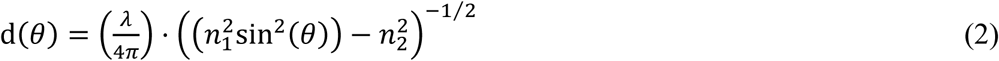

where *I*_0_(*θ*) is the maximum pixel intensity, d(*θ*) is the penetration depth of the evanescent field at an incident angle *θ*, *λ* is the wavelength of the illumination light, and *n*_1_ and *n*_2_ denote the refractive indices of the solution and glass coverslip, respectively.^37^ *I*_0_(*θ*) was assigned as the closest point to the coverslip, allowing us to generate a 3D membrane topography map. The generated images were thresholded (minimum intensity = 200) to create binary masks that isolated microvilli regions from the background. External contours of the microvilli regions were detected using an edge detection algorithm (Figure 3e). Using the height-profile images, we generated corresponding contour maps of the individual microvilli (Figures 3f, g).^33^

The scanning electron microscopy (SEM) image of the KG1a cells deposited on a coverslip showed a spherical shape of the cell with microvilli protruding structures (Figure 3h). The density of the microvilli identified by the TIRFM-based surface topography mapping (2.5 per μm^2^) is approximately half of that determined by SEM (5.0 per μm^2^) (Figure 3i), which is due to the different XY spatial resolution of the two imaging methods and the varied length of the microvilli. Nevertheless, the result suggests that the TIRFM-based cell surface topography mapping can relatively efficiently capture the 3D structure of the individual microvilli of the KG1a cells.

### 3D localization of adhesion molecules using SMLM and integration of single-molecule coordinates with 3D topographic maps

To achieve a nanoscale spatial correlation between the cell surface morphology captured by the TIRFM imaging and the 3D coordinates of the adhesion molecules characterized by the astigmatism-based 3D-SMLM imaging, we obtained TIRFM and 3D-SMLM data sequentially from the same KG1a cell under two different illumination conditions (i.e., TIR and HILO for TIRFM and 3D-SMLM imaging, respectively). Our custom-built microscopy setup allowed a reliable reversible switching between the two illumination modes (Figure S2). The cell membrane of the KG1a cells was fluorescently labeled by the MemBrite Fix 640, whereas the adhesion molecules were immunolabeled with Alexa Fluor 488-conjugated antibodies. After fixing, the cells were placed on a silane-coated glass surface in the switching buffer. We first acquired the TIRFM images of MemBrite Fix 640 using a 640 nm excitation laser, then acquired the 3D-SMLM images using a 488 nm laser. The 3D-SMLM images were captured precisely at the interface between the cell and the coverslip, where microvilli structures were most prominently observed (Figure 4a). The axial localization precision of our experiment was approximately 30 nm (Figure S6). The obtained single-molecule 3D coordinates of the adhesion molecules (CD44 in Figure 4a) were first filtered by the Z-axis range (i.e., the localizations that fall within the 120 nm from the coverslip surface, Figure 4b), then filtered based on their spatial position relative to the binary mask corresponding to the contour area of the microvilli (Figure 4c). Only molecules with XY coordinates falling within or on the boundary of a microvilli contour were retained. Then, the spatial relationship between microvilli topography and spatial coordinates of the adhesion molecules was visualized by overlaying the cell surface topography map with the single-molecule localizations of the adhesion molecules (Figure 4d).

**Figure 4.**
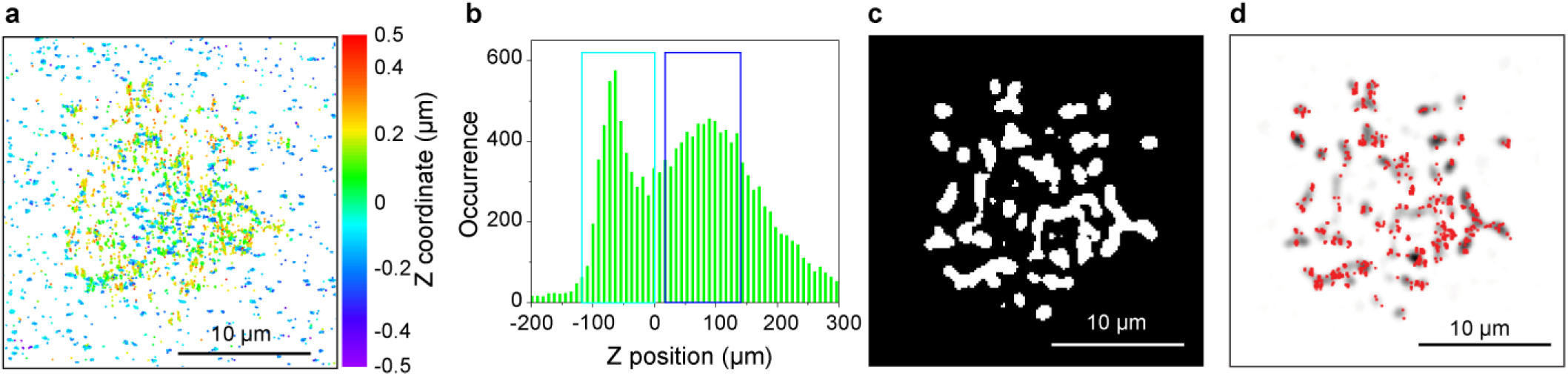
3D coordinates of adhesion molecules on microvilli determined by 3D-SMLM. Panels (a–d) illustrate the data processing pipeline that filters and overlays SM localizations relative to microvilli structures. (a) 3D-SMLM localization map of CD44 on a KG1a cell, immunolabeled with AF-488-conjugated antibodies, captured at 0.6 μm z-axis depth. The color bar represents the z-axis position. The 3D-SMLM image was obtained from the same cell shown in Figure 3. (b) Z-coordinate frequency distribution of individual localized CD44 molecules in (a). The cyan rectangle indicates the molecules adsorbed on the coverslip, whereas the blue rectangle highlights the molecules on the cell surface located within a 120 nm distance from the coverslip that were used for the following XY-filtration. (c) Binary mask image showing microvilli contour regions obtained from the 3D topographic map shown in Figure 3e with the threshold value of 200 counts. Mask applied to Z-filtered molecules, revealing that selected molecules localize to microvilli. (d) 3D overlay projection of the 3D topographic image obtained by TIRF microscopy displayed in the grayscale and the 3D coordinates of CD44 determined by 3D-SMLM and filtered by XY-filtration using the binary mask shown in red dots.

### Validation of the performance of the method

To validate the accuracy of the 3D spatial overlay of the cell surface topography map and the 3D coordinates of the adhesion molecules, we analyzed the spatial coordinates of actin molecules with respect to the cell surface topography map. Actin filaments are key cytoskeletal components that drive microvilli protrusion and contribute to their highly curved structure.^43^ Since their spatial localization within microvilli is well known, it is an ideal reference adhesion molecule for validating the accuracy of our method. The MemBrite Fix 640 stain was used for the TIRFM-based surface topography mapping, whereas Alexa Fluor 488-conjugated phalloidin was used to label the actin cytoskeleton. The overlaid topography map and 3D coordinates (Figure 5a) were segregated into multiple regions of interest (ROI), each containing a microvillus (Figure 5b). The 3D view of the overlaid image (Figure 5b) clearly showed that actin aligned within the microvillus structure, which is the expected spatial relationship between actin filaments and microvilli. The overlaid image captured the nanoscale three-dimensional spatial architecture more clearly than the XY projection of the localizations overlaid with the surface topography image that can be obtained using previously reported combined TIRF microscopy and 2D SMLM (Figure S7).^32, 34, 39^ The bivariate histograms generated using the 3D coordinates of the actin molecules clearly showed a filamentous shape of actin inside the microvillus (Figures 5c, d), further validating the accuracy of the spatial overlaying of the two 3D images.

**Figure 5.**
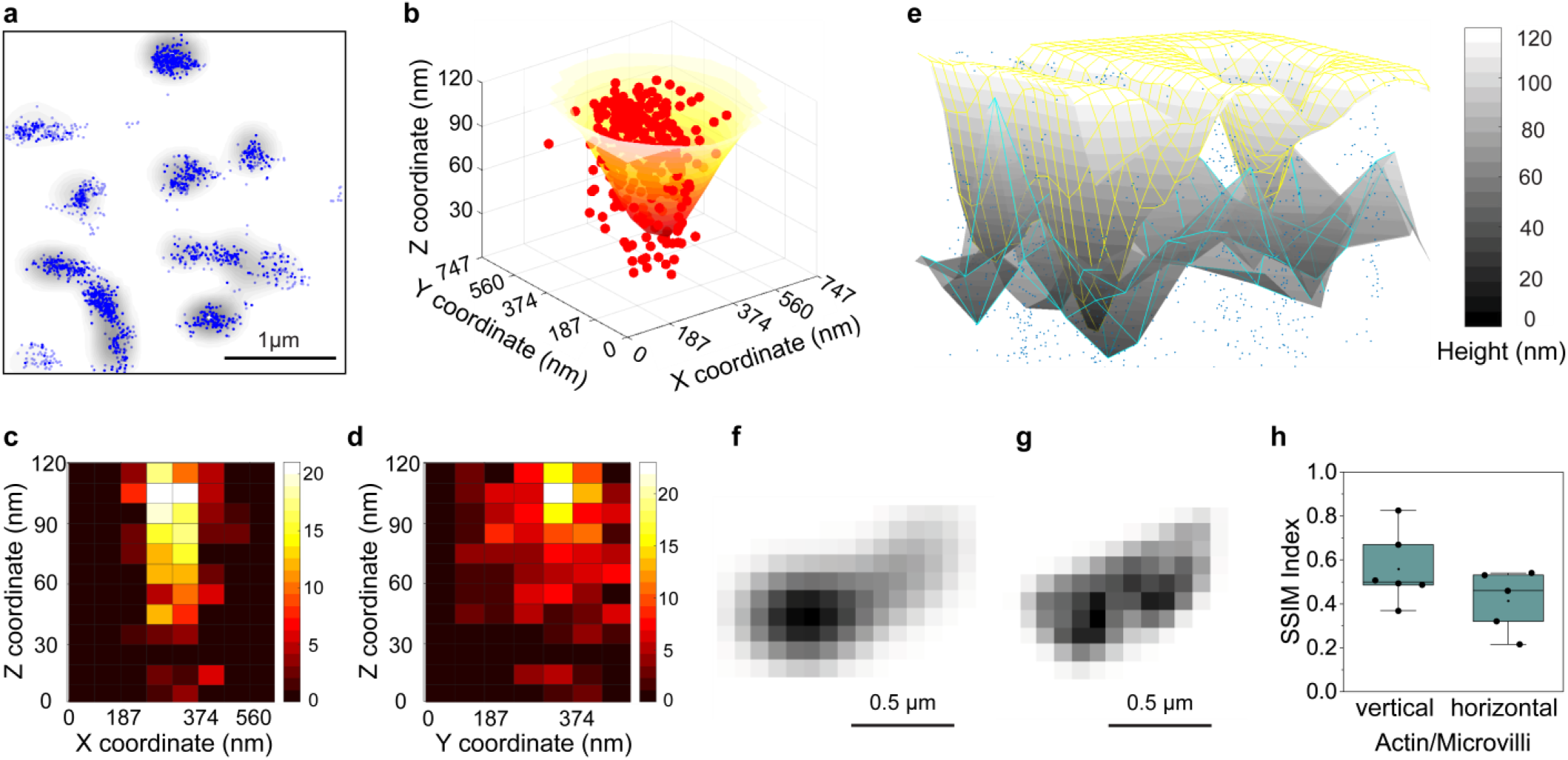
Evaluation of the spatial superposition of the 3D cell surface topography and 3D coordinates of adhesion molecules. (a) 3D-SMLM localization map of actin on a KG1a cell labeled by AF-488-conjugated phalloidin and overlaid with a 3D cell surface topography image acquired by TIRFM with MemBrite FX640 stain. Blue dots indicate localized individual actin molecules, and the grayscale image shows the contours of microvilli on the cell surface. (b) 3D view of localized individual actin molecules superimposed on a single microvillus on a KG1a cell. (c, d) Bivariate frequency histograms showing the 3D distribution of actin molecules in the microvillus on a KG1a cell, projected along the (c) XZ and (d) YZ axes. (e) 3D view of the cell surface topography shown as a yellow mesh, overlaid with the spatial boundary obtained from the 3D coordinates of individual actin molecules shown as a cyan mesh. Blue dots indicate localized individual actin molecules. The axial distance from the coverslip surface is shown in grayscale. (f, g) Normalized 8-bit 2D images obtained from the (f) 3D topography of a single microvillus and (g) 3D spatial boundary obtained from the 3D coordinates of individual actin molecules. The axial distance from the coverslip surface is shown in grayscale. (h) Structural similarity (SSIM) index analysis of the 3D topography of the vertically- and horizontally-oriented microvilli generated by TIRFM and the 3D coordinates of the actin molecules determined by 3D-SMLM.

We quantified the overlaying performance by calculating the structural similarity (SSIM) index between the 3D surface topography map obtained using the TIRFM and the spatial boundary obtained by 3D surface fitting of the 3D coordinates of the actin molecules (Figure 5e). The SSIM index values were calculated using the normalized topography map and the normalized spatial boundary by converting them into 2D grayscale images (Figures 5f, g, h, and S8a, b and S9a), preserving the spatial structure of the 3D volume (see Materials and Methods for the details). The mean SSIM index value (SSIM_mean_ = 0.50, Figure S9a) suggests a high spatial overlapping of the two images, confirming the good overlaying performance of our method. In addition, the SSIM analysis showed that the vertically oriented microvilli (Figure 5h and S8a) gave a larger SSIM index (SSIM_mean_ = 0.56) than the horizontally oriented microvilli ((SSIM_mean_ = 0.41, Figures 5h and S8b). This result suggests that the TIRF-based reconstruction of the 3D topography of the microvilli worked better for the vertically-oriented microvilli, indicating a potential artefacts in the reconstructed 3D topography of the horizontally-oriented microvilli. This is expected as the conversion of the TIRF intensity into the topographic map assumes the cell membrane at any XY position has a single axial position, which is not the case for the horizontally-oriented microvilli that would have two axial positions at the same XY position (i.e., top and bottom membranes of the microvilli). These results suggest that our data analysis pipeline enables accurately localizing the 3D coordinates of the adhesion molecules on the microvilli without artefacts.

### Spatial distribution of membrane adhesion proteins in relation to the 3D topography of the microvilli

Having demonstrated the effectiveness of our combined TIRFM and 3D-SMLM in mapping the actin cytoskeleton relative to microvilli, we next applied our method to other adhesion molecules. Specifically, we focused on key adhesion molecules in the initial homing step of HSPCs, CD44 and PSGL-1, which facilitate tethering and rolling of HSPCs. Our previous studies have revealed their distinct colocalization patterns, with CD44 being highly expressed and homogeneously distributed across the cell membrane, whereas PSGL-1 exhibits an inhomogeneous distribution.^8,9^ Immunogold labeling studies have further indicated that PSGL-1 preferentially localizes to microvilli, while CD44 is evenly distributed across the cell body.^44,45^

We performed the imaging experiments on fixed KG1a cells mounted on a silane-coated glass coverslip that were stained by the MemBrite Fix 640 stain and immunolabeled with Alexa 488-conjugated antibodies targeting CD44 and PSGL-1. The 3D views of the microvilli showed spatial colocalization of microvilli with CD44 (Figure 6a) and PSGL-1 (Figure 6e). While both adhesion molecules localize on the microvilli, the 3D views clearly revealed distinct spatial patterns of these adhesion molecules. The CD44 molecules are distributed almost uniformly on the microvilli (Figure 6a), whereas the PSGL-1 molecules show a non-random distribution on the microvilli (Figure 6e). The bivariate histograms generated using the 3D coordinates of the CD44 molecules clearly captured their distribution on the surface of the microvilli (Figures 6b, c). On the other hand, the bivariate histograms of the PSGL-1 molecules revealed the formation of clusters on the microvilli (Figures 6f, g). Overall, the spatial distributions of CD44 and PSGL-1 on the microvilli obtained in this study are consistent with those indicated by previous studies. Interestingly, our new method revealed that the PSGL-1 clusters are located not only at the tip of the microvilli, as indicated by previous studies,^46,47^ but also on other parts of the microvilli. The comparison with the XY projections of the 3D images (Figures S7b, c) demonstrates that the 3D nanoscopic architectures of the adhesion molecules that cannot be resolved in the 2D projection are clearly captured using our new method. The mean SSIM index values of CD44/microvilli (SSIM_mean_ = 0.44) and PSGL-1/microvilli (SSIM_mean_ = 0.55) confirm overall good spatial overlay of the 3D topographic image of the microvilli and 3D localizations of the adhesion molecules (Figures S9b, c), reassuring the reliable spatial overlay of the two images in our method. The SSIM analysis also showed that the vertically oriented microvilli gave larger SSIM index (SSIM_mean_ = 0.46 and 0.58 for CD44 and PSGL-1, respectively) than the horizontally oriented microvilli (SSIM_mean_ = 0.39 and 0.38 for CD44 and PSGL-1, respectively, Figures 6d, h), similar to the actin molecules, indicating the importance of selecting only the vertically-oriented microvilli for the analysis without artifacts.

**Figure 6.**
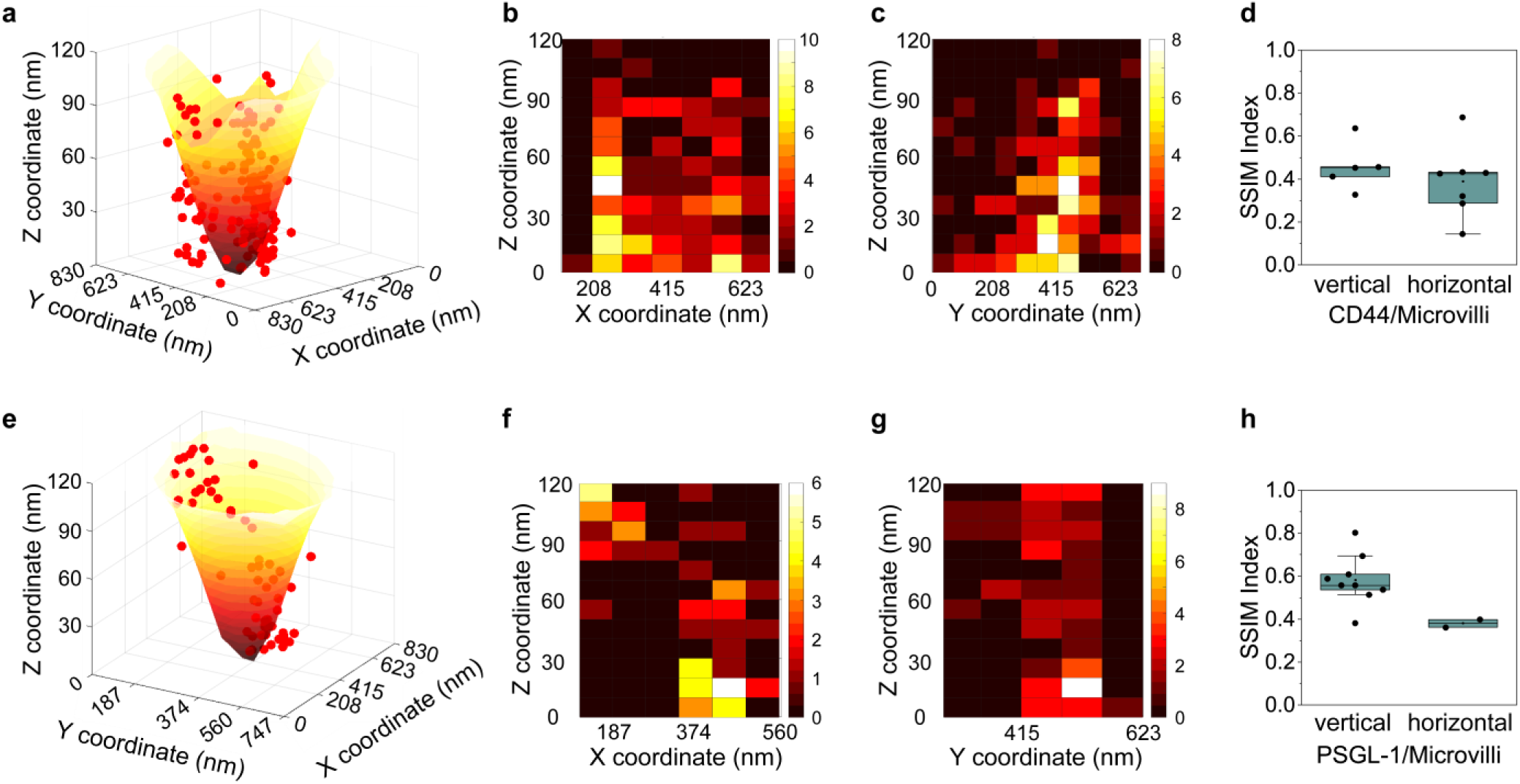
Spatial distribution of adhesion molecules on microvilli. (a) 3D view of localized individual CD44 molecules immunolabeled with AF-488-conjugated antibodies, overlaid onto a single microvillus on a KG1a cell. (b, c) Bivariate frequency histograms showing the 3D distribution of CD44 molecules on the microvilli on a KG1a cell, projected along the (b) XZ and (c) YZ axes. (d) Structural similarity (SSIM) index analysis of the 3D topography of the vertically- and horizontally-oriented microvilli generated by TIRFM and the 3D coordinates of the CD44 molecules determined by 3D-SMLM. (e) 3D view of localized individual PSGL-1 molecules immunolabeled with AF-488-conjugated antibodies, overlaid onto a single microvillus on a KG1a cell. (f, g) Bivariate frequency histograms showing the 3D distribution of PSGL-1 molecules on the microvillus on a KG1a cell, projected along the (f) XZ and (g) YZ axes. (h) SSIM index analysis of the 3D topography of the vertically- and horizontally-oriented microvilli generated by TIRFM and the 3D coordinates of the PSGL-1 molecules determined by 3D-SMLM.

### Characterization of the effect of cell rolling on the spatial distribution of membrane adhesion proteins

Our previous studies demonstrated the spatial reorganization of CD44 and PSGL-1 during cell rolling over E-selectin, which is associated with the nanoscopic morphology change of the cell membrane, including the formation of the membrane tethers and slings, caused by the shear stress exerted on the rolling cells.^8,9,28^ Thus, we next examined the applicability of our method to the KG1a cells undergoing rolling over E-selectin. To that end, we integrated the TIRFM and 3D-SMLM imaging setup into a microfluidics-based cell rolling assay platform (Figure 7a).

**Figure 7.**
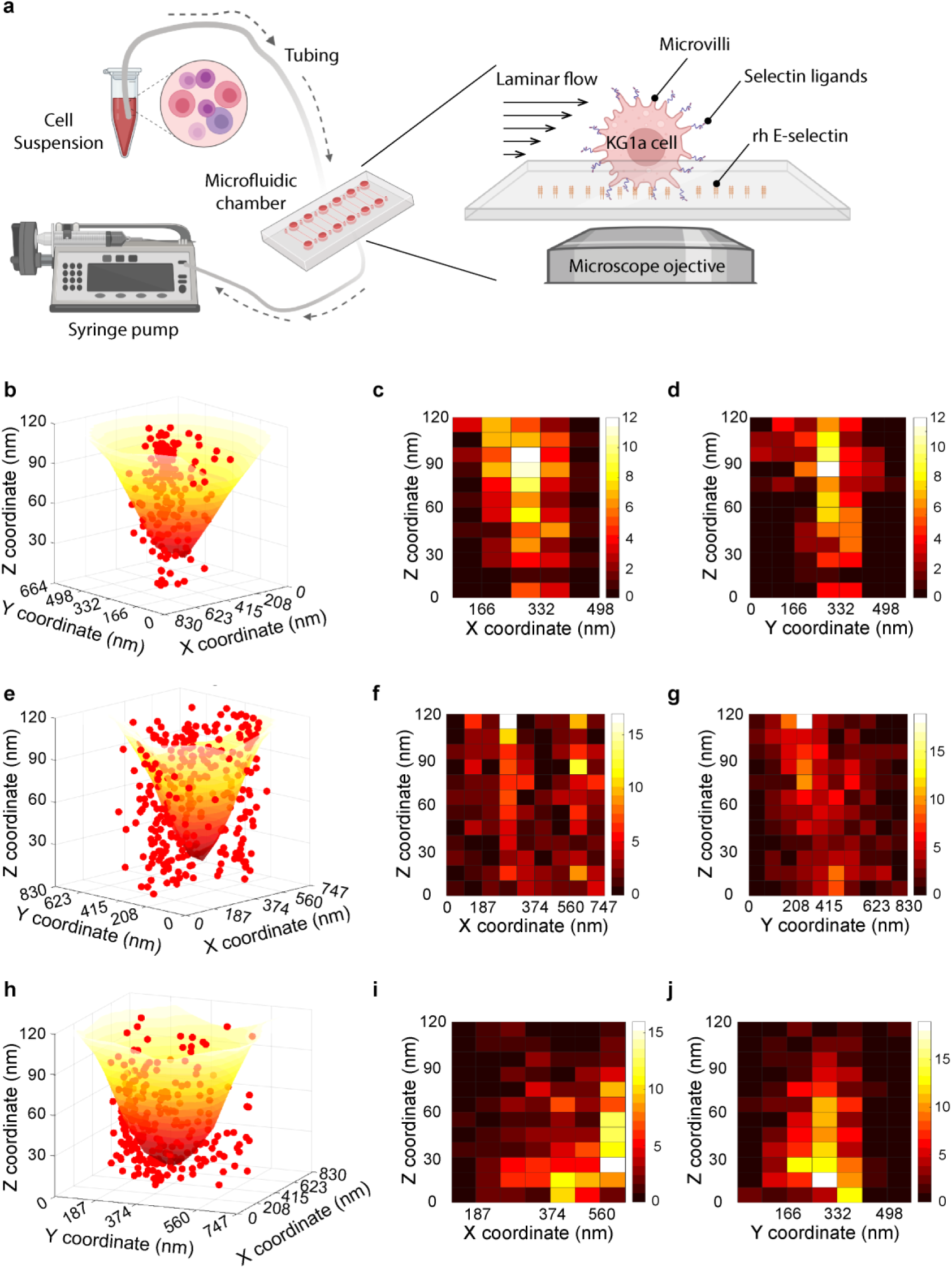
Effect of cell rolling on the nanoscopic 3D distribution of adhesion molecules relative to the 3D Microvilli topography. (a) Schematic illustration of the cell rolling assay using a microfluidic device. A suspension of KG1a cells was infused into the fluidic chamber coated with rh E-selectin using a syringe pump. (b) 3D view of localized individual actin molecules superimposed on a single microvillus on a KG1a cell rolled over E-selectin. (c, d) Bivariate frequency histograms showing the 3D distribution of actin molecules in the microvillus on a KG1a cell rolled over E-selectin, projected along the (c) XZ and (d) YZ axes. (e) 3D view of localized individual CD44 molecules immunolabeled with AF-488-conjugated antibodies, overlaid onto a single microvillus on a KG1a cell rolled over E-selectin. (f, g) Bivariate frequency histograms showing the 3D distribution of CD44 molecules on the microvilli on a KG1a cell rolled over E-selectin, projected along the (f) XZ and (g) YZ axes. (h) 3D view of localized individual PSGL-1 molecules immunolabeled with AF-488-conjugated antibodies, overlaid onto a single microvillus on a KG1a cell rolled over E-selectin. (i, j) Bivariate frequency histograms showing the 3D distribution of PSGL-1 molecules on the microvillus on a KG1a cell rolled over E-selectin, projected along the (i) XZ and (j) YZ axes.

KG1a cells were first stained by the MemBrite Fix 640 stain, then perfused into the microfluidic chamber coated with recombinant human E-selectin (rhE-selectin) under physiological shear stress (2 dyne cm⁻²) using a syringe pump. After one minute of rolling, the KG1a cells were fixed in situ by perfusing a fixative solution and then immunolabeled against the target adhesion molecules before imaging under conditions identical to the TIRFM and 3D-SMLM imaging experiments on the controlled cells (i.e., KG1a cells rested on the glass coverslip, see Materials and Methods section for the details of the experimental procedures). The obtained TIRFM and 3D-SMLM data were analyzed using the image processing pipeline identical to that applied to the control cells.

The overlaid topographic image of the microvilli and the 3D localizations of the adhesion molecules showed that the spatial distributions of the actin and CD44 molecules on the microvilli were not affected significantly during the rolling of the KG1a cells on E-selectin (Figures S10 and S11). The images also showed an increase in the footprint upon cell rolling (i.e., larger surface areas are in close proximity to the E-selectin-coated surface). This observation is consistent with our previous study, which reported the increased footprint of CD44 upon cell rolling, characterized by 2D-SMLM.^8^ A good spatial overlap between the microvilli and the actin and CD44 molecules after cell rolling is also consistent with our previous study using 2D-SMLM.^8^ The 3D view of the microvillus overlaid with the actin molecules (Figure 7b) and their bivariate frequency histograms (Figures 7c, d) further confirm that the spatial architecture of the actin cytoskeleton inside the microvillus is maintained during cell rolling. This result also demonstrates that the combined TIFRM and 3D-SMLM we developed in this study can report the accurate 3D structure of the microvilli and 3D localizations of the adhesion molecules on the microvilli, even under the flow condition using the microfluidic device. The actin filaments maintained an even distribution upon cell rolling (Figures 7b, c, d), indicating potential actin cross-linkage with adhesion proteins.^48,49^ The 3D view of the microvilli overlaid with the CD44 molecules (Figure 7e) and their bivariate frequency histograms (Figures 7f, g) clearly showed that the CD44 molecules distribute almost uniformly on the entire microvilli, suggesting their spatial architecture is maintained during cell rolling.

In contrast, PSGL-1 exhibited a large change in its spatial architecture on the microvilli upon cell rolling. The overlaid topographic image of the microvilli and the 3D localizations of the PSGL-1 molecules showed that PSGL-1 became more abundant and exhibited higher localization density upon cell rolling (Figure S12). The 3D view of the microvilli overlaid with the PSGL-1 molecules (Figure 7h) and their bivariate frequency histograms (Figures 7i, j) show the localization of PSGL-1 at the tip of microvilli, indicating that cell rolling affects the spatial architecture of PSGL-1 on the microvilli and promotes the spatial clustering of PSGL-1 at the microvilli tip. The XY projections of the 3D images (Figures S13) do not clearly reveal the reorganization of the adhesion molecules upon cell rolling, demonstrating the advantages of capturing 3D images using our new method. The observed nanoscopic reorganization of the spatial architecture of PSGL-1 demonstrates the capability of our new method in capturing the nanoscale spatiotemporal behavior of the cell surface morphology and the adhesion molecules in 3D.

## CONCLUSIONS

Microvilli, finger-like projections enriched with adhesion molecules, play a crucial role in HSPC homing by serving as anchoring and tethering points. Traditional models often overlook the 3D structure of the plasma membrane, instead treating it as a flat surface, which limits our understanding of adhesion molecule distribution.^50-55^ In this study, we reconstructed the 3D topography of KG1a microvilli and mapped key adhesion molecules by developing a new experimental approach, which integrates two high-resolution optical microscopy techniques, TIRFM and 3D-SMLM. Our method enabled precise 3D localization of adhesion molecules up to 500 nm along the Z-axis with 30 nm resolution and overlaid them to the microvilli on the cell surface.^32, 33, 42, 56^ Despite the diffraction-limited lateral resolution of TIRFM, our data showed that over 50% of the microvilli can still be reliably captured, demonstrating its effectiveness in spatially resolving cell surface structures.

We addressed potential imaging artifacts by employing SSIM analysis, which validated the accurate overlay of the TIRFM and 3D-SMLM datasets while mitigating signal amplification at microvilli tips. Our findings confirmed that the selectin ligands, CD44 and PSGL-1, pre-clustered within microvilli, forming distinct spatial organization patterns before and after rolling. Notably, PSGL-1 exhibited tip enrichment, whereas CD44 displayed a more uniform distribution, indicating that selectin/ligand interactions are structurally regulated at the nanoscale level. This spatial organization could enhance ligand exposure to selectins, increasing binding efficiency and stabilizing rolling adhesion. These results complement earlier work from our group showing that refining homing-specific adhesion mechanisms can significantly improve HSPC migration and engraftment ^4,7^, that engineered α1,3-fucosylation enhances selectin binding^57^, and that structural features beyond the lectin domain govern E-selectin–ligand affinity^24^. Collectively, our findings highlight microvilli as key signaling hubs for selectin-ligand interactions, emphasizing their role in the initial steps of homing.^4, 6, 7, 9, 51, 52, 57-60^

Beyond its application to studying HSPC and leukemic cell rolling, this combined TIRFM and 3D-SMLM approach can be broadly applied to investigate nanoscale spatiotemporal dynamics in a variety of biological systems. The ability to simultaneously resolve cell surface topography and molecular localization in three dimensions under physiologically relevant flow conditions makes it suitable for studying other adhesion-mediated processes, including immune cell trafficking, cancer cell metastasis, and pathogen-host interactions. In addition, because the method can be adapted to different molecular targets and experimental environments, it offers a versatile platform for uncovering how nanoscale membrane organization influences cellular behavior across diverse physiological and disease contexts.^61^

Moving forward, advancements in artificial intelligence-powered super-resolution techniques, such as Deep-STORM^62,63^ and ANNA-PALM,^64^ may further improve spatial-temporal resolution, allowing for real-time tracking of microvilli dynamics and selectin clustering.^63-70^ Future studies should aim to integrate these AI-based imaging approaches to capture real-time molecular reorganization during HSPC rolling and adhesion. Finally, our developed method is versatile and applicable to various in vivo and in vitro models, enabling nanoscale characterization of biological phenomena.

## MATERIALS AND METHODS

### Cells and treatments

KG1a cells, a human acute myelogenous leukemia cell line (ATCC), were maintained in RPMI (Gibco) supplemented with 10% fetal bovine serum, penicillin (100 U ml^-1^), and streptomycin (100 µg ml^-1^). Cells in suspension were maintained at 37°C in a humidified atmosphere containing 5% CO2. The viability of the cells was routinely checked by trypan blue staining or calcein AM dye. The cell culture medium was refreshed a day before preparing the experimental samples.

### Microfluidic chamber preparation

Glass coverslips (No. 1.5, ibidi GmbH) were cleaned by ultrasonication (P60H, Elma Schmidbauer GmbH) in potassium hydroxide and ethanol. A clean coverslip was attached to the bottom of a microfluidic chamber (channel width, 3.8 mm; channel height, 0.4 mm; sticky-Slide VI 0.4, ibidi GmbH) and incubated with protein A (10 μg ml^−1^; Invitrogen) overnight at 4°C. After washing off unbound protein A with HBSS, the chamber was incubated with a rhE-selectin (Sino Biological), which contained the fused C-terminal polyhistidine-tagged Fc region of human immunoglobulin G1 (IgG1) at the C terminus, at 4°C at concentrations of 0.2 μg ml^−1^ for 1 hour. The chamber was then washed with HBSS and blocked with 1% casein solution in phosphate-buffered saline (PBS; Thermo) for 30 min to 1 hour at room temperature. The E-selectin deposited chamber was used immediately for the cell-rolling assay and SR imaging experiments.

A clean glass coverslip was coated with silane before attaching it to the bottom of the microfluidic chamber to prepare a coated surface for control samples. Silane coating was performed following the manufacturer’s instructions. First, traces of water were removed from the clean coverslip using acetone. Second, silane treatment was performed by soaking the dehydrated coverslips in 2% (v/v) silane solution by mixing 3-aminopropyltriethoxysilane (Sigma A3648) in acetone for 2 minutes. To quench the silanization reaction, the coverslip was soaked in water (dH2O), which allowed for the rapid exchange of the silane solution with water. Before attaching the silane-coated coverslip to the bottom of the microfluidic chamber, the coverslip was placed into the oven and baked for 30 minutes at 110 degrees to cure the silane.

### Cell-rolling assay

The cell-rolling assay was performed at room temperature using a microfluidic chamber sticky-Slide VI 0.4 attached to the E-selectin–deposited glass coverslip. The inlet and outlet of the chamber were connected to a 0.8-mm silicon tubing (ibidi GmbH) using male Luer connectors (ibidi GmbH). The end of the inlet tube was placed in a rolling buffer [HBSS containing 1% human serum albumin (Sigma) and 1 mM CaCl_2_ (Sigma)], and the end of the outlet tube was connected to a programmable syringe pump (PHD ULTRA, Harvard Apparatus) using a female Luer Lock connector (ibidi GmbH). 10^6^ ml^−1^ of MemBrite-stained KG1a cells suspended in the rolling buffer were perfused inside the chamber. After a brief stop in flow for 1 min to allow the cells to settle down and interact with the E-selectins on the surface, the chamber was washed with the rolling buffer for 90 to 120 seconds, during which we observed the cells rolling over the E-selectin–coated surface of the chamber. The rolling experiment was conducted at wall shear stresses (W) of 2 dyne cm^−2^. The cell-rolling behavior was observed by mounting the microfluidic chamber on an inverted optical microscope (CXK41, Olympus) equipped with a 20× objective (LCAch N 20X, Olympus) and a CCD camera (XC10, Olympus) using CellSens software (Olympus). To capture immunofluorescence images of rolled cells, a fixation solution of 4% paraformaldehyde and 0.2% glutaraldehyde was perfused into the microfluidic chamber and incubated for 15 minutes at room temperature. Then, the cells were washed with a buffer for 5 minutes at room temperature and utilized for fluorescence immunolabeling.

### Fluorescence labeling of the cells

A suspension of KG1a cells was washed with prechilled 1X HBSS for 5 min and centrifuged at 4°C. Immunostaining of the KG1a cells for two-color fluorescence imaging was conducted following the manufacturer’s recommendation. First, live KG1a (10^6^ ml^−1^) cells were incubated in 2X pre-staining solution in HBSS buffer for 10 min at 37°C. Then, the pre-staining solution was removed, and the cells’ membranes were stained with 2X Membrite Fix640 (Biotium) solution in HBSS buffer for 30 min at 4°C. Then, the cells were either directly fixed with a fixative buffer of 4% (w/v) paraformaldehyde (Electron Microscopy Sciences) and 0.2% (w/v) glutaraldehyde (Electron Microscopy Sciences) in HBSS for 10 min at room temperature or utilized for the rolling experiment, then fixed. After fixing the control and rolling samples, cells were washed gently 2 times and blocked using 10% goat serum (Sigma) for 40 min at RT. The cells were incubated with purified mouse anti-human CD44 antibody (15 μg ml^−1^; clone 515, BD Pharmingen), mouse anti-human PSGL-1 antibody (15 μg ml^−1^; clone KPL-1, BioLegend), diluted in 2% BSA (Sigma) in HBSS for 40 min at room temperature, followed by AF-488–conjugated goat anti-mouse secondary antibody (5 μg ml^−1^; Invitrogen) diluted in 2% BSA in HBSS for 40 min at room temperature. Then, the cells were fixed again in 3% (w/v) paraformaldehyde and 0.2% (w/v) glutaraldehyde in HBSS for 10 min at room temperature. Immobilized immunolabeled KG1a cells were stored in PBS buffer overnight at 4°C. Before performing SR imaging, a freshly prepared imaging buffer was perfused gently into the chamber to replace the HBSS buffer. Control cells were suspended in an imaging buffer before perfusing them inside a silane-coated microfluidic chamber.

For actin cytoskeleton labeling, the membrane-stained and fixed cells were then permeabilized in 0.1% Triton X-100 (Sigma-Aldrich) in cytoskeleton buffer [10 mM MES (pH 6.1), 150 mM NaCl, 5 mM EGTA, 5 mM glucose, and 5 mM MgCl_2_] for 10 min at room temperature. The permeabilized cells were labeled for actin cytoskeleton with freshly prepared 0.5 μM AF-488 phalloidin (Molecular Probes-Thermo Fisher Scientific) diluted from a stock solution of 6.6 μM AF-488 phalloidin, with 1% BSA in the cytoskeleton buffer for 1 hour at room temperature. Then, the cells were fixed again in 3% (w/v) paraformaldehyde and 0.2% (w/v) glutaraldehyde in HBSS for 10 min at room temperature.

### Fluorescence microscopy

SR fluorescence imaging experiments were conducted using a home-built fluorescence microscopy setup (Figure S2).^71,72^ Briefly, a CW solid-state laser operating at either 640 nm or 488 nm (60 mW, Cobolt, MLD) that passed through an excitation filter (Semrock, LD01-640/8, or FF01-488/6 for the 488 nm excitation, respectively) and a beam expander 7X (Thorlabs) was introduced to the inverted microscope (Olympus, IX71) from its backside port through an achromatic convex lens (f = 300 mm; Thorlabs). The samples were illuminated using a TIRF or low-angle configuration through a high NA objective lens (Olympus, 100× NA = 1.49, oil immersion). Two-color fluorescence imaging experiments were conducted sequentially by introducing 640-nm and 488-nm lines of the lasers coaxially into the microscope’s rear port through a light path that adjusts the off-axis position of the focused laser beam at the back focal plane. The fluorescence from the sample was captured by the same objective, separated from the illumination light by a multiband dichroic mirror (Semrock, Di03-405/488/561/635-t1-25 × 36), and passed through a TuCam dual-camera adaptor (Andor Technology) equipped with a filter cassette containing a dichroic mirror (Semrock, FF580-FDi01-25 × 36) to separate the fluorescence into two channels. Two EMCCD cameras (Andor Technology, iXon3 897) detected the separated fluorescence from the samples through emission bandpass filters (Semrock, FF01-550/88-25 and FF01-697/58-25). The acoustic-optic tunable filter (AOTF), a laser control system (Andor Technology, PCUB-110), was utilized to synchronize the exposure time of the EMCCD camera and the illumination of the sample by the excitation laser. The image acquisition was performed using the Andor iQ3 software. Our setup had a final magnification of 192x, which resulted in a pixel size of 83.33 nm.

### Total internal reflection fluorescence microscopy (TIRFM)

Microfluidic-based TIRFM imaging was utilized to map the surface morphology of the cell. To achieve total internal reflection at the sample, the position of the focused beam was shifted from the center of the objective lens to its edge. The light path was comprised of a 90° reflective plane mirror and the achromatic convex lens (f = 300 mm; Thorlabs) mounted on the linear piezoelectric actuator stage (Thorlabs) that was controlled by Kinesis software (Thorlabs). The drive acceleration was set to 2 mm/s. The refraction index of the glass coverslip was *n*_e_ = 1.52, imaging buffer *n*_e_ = 1.33, while that of the immersion oil objective was *n*_e_ = 1.518. The bottom surface of the membrane-stained KG1a cell was illuminated at TIR angles 66.6° (2.59 mm away from the objective lens center off-axis position, which generated an evanescence wave with a penetration depth of 120 nm) under weak illumination of 640-nm laser power (equivalent to 30 W cm^−2^ at Epi configuration). Per a sample field of view, 10 frames were captured to average the fluorescence intensity before processing the TIRF image for the topographical map construction.

### 3D single-molecule localization microscopy (3D-SMLM)

3D-SMLM was utilized to capture the 3D locations of selectin ligands on the cell surface. 3D-SMLM imaging of immunolabeled KG1a cells was performed in an imaging buffer containing TN buffer [50 mM tris (pH 8.0) and 10 mM NaCl], oxygen scavenging system [glucose oxidase (0.5 mg ml^−1^; Sigma), catalase (40 µg ml^−1^; Sigma), 10% (w/v) glucose], and 10 mM Cystamine hydrochloride (pH 8.0) (Sigma) as a reducing reagent. The imaging solution was prepared immediately before the imaging experiments. The samples were illuminated using a low-angle-configured objective lens where the laser beam’s off-axis position was positioned 2.3 mm away from the objective lens center. The illumination power at the samples for the imaging experiments was set to 2 kW cm^−2^. The illumination area was adjusted to a diameter of 25 µm so that only a single cell was illuminated during each image acquisition. The fluorescence from the sample was captured by the same objective, separated from the illumination light by a dichroic mirror (FF660-Di02-25×36, Semrock), and detected by an EMCCD camera after passing through an emission bandpass filter (FF01-697/58-25, Semrock). The fluorescence images were captured using a 200 Å∼200-pixel region of the EMCCD camera with 83-nm pixel size and a 30-ms exposure time. 10,000 fluorescence image sequences were captured to reconstruct SR localization microscopy images. The image acquisition was utilized using the Andor iQ3 software. The exposure of the EMCCD camera was synchronized with the sample illumination by the laser using an AOTF (AA Opto Electronic).

The 3D-SMLM imaging experiment was conducted using the illumination configuration and buffer condition in the same manner as the 2D imaging experiment. Astigmatism-based SR localization microscopy was implemented for the 3D SR imaging.^73^ A 1000 mm focal length cylindrical lens was inserted before the imaging lens. The z-axis positions were calibrated using 20 nm-diameter FluoSpheresTM carboxylate, yellow-green (505/515) (Invitrogen) deposited on a cleaned coverslip in HBSS buffer 1.0 x 10^-8^ dilution. The calibration data was collected with a 10-nm step size between −300 and +300 nm. A piezo nanopositioning stage (APZ-X100 Piezo Z-Stage, Andor Technology) controlled the z-axis positions during data acquisition for calibration.

The localization precision SD of the Gaussian = 30 nm. The calibration data were fitted to the elliptical 2D Gaussian function, and the z-position-dependent spot widths were utilized to obtain the calibration curve. The stage drift in the z-axis was less than 30 nm during each image acquisition (∼180 s), and it was stabilized at a constant focal plane using the C-focus system (MCL). During the SR image acquisition, the offset was set between -0.5 and +0.5 mV.

### Analysis of the 3D-SMLM images

The 3D-SMLM images were reconstructed using a custom-written MATLAB (MathWorks) code or Localizer software.^74^ The positions of the AF-488 molecules were determined by fitting the particle localization with a 2D Ellipsoidal Gaussian fitting (astigmatism) function. Fluorescence spots whose width was significantly larger (>200 nm) than the PSF of the optical system (PSF, ∼130 nm) were excluded. Stage drift was corrected by reconstructing the sub-images using 5000 localizations. The calibration curve was obtained in the 3D-SMLM imaging by fitting the calibration data to the elliptical 2D Ellipsoidal Gaussian fitting (astigmatism) function and fitting the z–position–dependent spot widths to polynomial functions. TetraSpeck microspheres (diameter, 100 nm) deposited on a cleaned coverslip were used to calibrate the shift between the two channels in the two-color SR imaging. Using fluorescence images of the TetraSpeck microspheres captured simultaneously by two cameras, we generated a registration map that corrected the shift between them and applied the correction to the cell sample images.

### Scanning electron microscopy

KG1a cells resting on a glass slide were prepared using a previously published protocol.^9^ The harvested KG1a cells (107 cells) were washed twice with HBSS. The cells were fixed in 2-2.5% glutaraldehyde in 0.1 M Cacodylate buffer (pH = 7.2–7.4) at 4 °C overnight. The fixed cells were washed three times with the 0.1 M Cacodylate buffer, submerged for 15 min in the buffer before the next wash, and resuspended in 200 μl of the buffer.

Then, 100 μl of the cell suspension was added to a cover slip, which was brushed with polylysine and incubated overnight inside a moisturized chamber before use. The cells were further fixed in 1% osmium tetroxide in 0.1 M Cacodylate buffer for one hour in the dark. The cells were washed three times with distilled water, where they were submerged in water for 15 min before the next wash, and dehydrated in gradient ethanol (30, 50, 70, 90, and 100%). Samples were carried further onto the Critical Point Drying apparatus and covered with 100% ethanol to submerge the samples completely. The dehydrated cells were placed on a holder of the scanning electron microscope (SEM) and coated with 4 mm platinum (K575X Sputter Coater, Quorum). The SEM images were recorded using Teneo SEM (Thermo Fisher).

### Structural similarity (SSIM) index analysis

The structural similarity (SSIM) index between the 3D surface topography map obtained using the TIRFM and the spatial boundary obtained by 3D surface fitting of the 3D coordinates of adhesion molecules. The SSIM index values were calculated using the normalized topography map and the normalized spatial boundary by converting them into 8-bit 2D grayscale images with the same pixel size and pixel number. Then, the SSIM index was calculated pixel-wise.

### Statistics

The student’s t-test assessed statistical significance (assuming two-tailed distribution and two-sample unequal variance). All experiments were repeated at least three times to ensure reproducibility. All of the single-cell fluorescence microscopy images reported in this study are representative examples of multiple (n > 3) independent experiments.

## ASSOCIATED CONTENT

### Supporting Information

The following files are available free of charge. Supporting texts describing the calculation of the critical angle of the total internal reflection and the maximum angle in TIRFM, and verification of the incident angles and penetration depths of the excitation light, Schematic overview of the hematopoietic stem/progenitor cell homing mechanisms (Figure S1), Schematic diagram of a combined TIRF and 3D-SMLM setup (Figure S2), Uniform labeling of the cell membrane using membrane stains (Figure S3), Determination of the incident angles (Figure S4), Verification of the incident angles and penetration depths of the excitation light (Figure S5), Localization precision of 3D-SMLM (Figure S6), Spatial distribution of adhesion molecules on microvilli projected on the XY plane (Figure S7), Comparison of the 3D topographic image of the cell surface and the 3D spatial boundary obtained from the 3D coordinates of individual actin molecules (Figure S8), Structural similarity index analysis of the 3D topography of the microvilli generated by TIRFM and the 3D coordinates of the adhesion molecules determined by 3D-SMLM (Figure S9), Effect of cell rolling on the architecture of microvilli and spatial distribution of actin (Figure S10), Effect of cell rolling on the architecture of microvilli and spatial distribution of CD44 (Figure S11), Effect of cell rolling on the architecture of microvilli and spatial distribution of PSGL-1 (Figure S12), Spatial distribution of adhesion molecules on microvilli on the rolled cells projected on the XY plane (Figure S13) (PDF)

## AUTHOR INFORMATION

### Corresponding Author

\***Satoshi Habuchi** - Biological and Environmental Science and Engineering Division, King Abdullah University of Science and Technology, Thuwal 23955-6900, Saudi Arabia

### Author Contributions

The manuscript was written through the contributions of all authors. All authors have given approval to the final version of the manuscript. †These authors contributed equally.

## ACKNOWLEDGMENT

This study was supported by King Abdullah University of Science and Technology (KAUST).

