## Supporting Information for "Combined TIRF and 3D Super-Resolution Microscopy for Nanoscopic Spatiotemporal Characterization of Adhesion Molecules on Microvilli"

**Contents**

Supporting Texts 1 to 2

Supporting Figures 1 to 13

### Supporting Text

#### 1. Calculation of the critical angle of the total internal reflection (TIR) and the maximum angle in TIRFM

The critical angle at which total internal reflection occurs ($\theta_{\mathrm{critical}}$) at the glass/buffer interface was determined using the following equation:

$$\theta_{\mathrm{critical}} = \sin^{-1}\left(\frac{n_1}{n_2}\right) = 61.04°$$

where $n_1$ = 1.33 (cell cytosol) and $n_2$ = 1.52 (refractive index of the coverglass and immersion liquid). The maximum achievable TIR angle ($\theta_{\mathrm{max}}$), constrained by the numerical aperture (NA) of the TIRF objective lens (NA = 1.49), was calculated using the following equation:

$$\sin\theta_{\mathrm{max}} = \frac{\mathrm{NA}}{n_2} = 78.6°$$

#### 2. Verification of the incident angles and penetration depths of the excitation light

To verify the accuracy of the incident angles of the TIR illumination and, therefore, the penetration depth of the evanescent field, we measured integrated fluorescence intensities under varied TIR illumination conditions (i.e., varied TIR angles). The integrated fluorescence intensity ($I$) obtained from a solution containing a uniform concentration of fluorophores illuminated at a specific TIR angle ($\theta$) is obtained using equations 1 and 2 and is described as follows:

$$I = I_0\left(\frac{\lambda}{4\pi\sqrt{n_1^2\sin^2\theta - n_2^2}}\right) \qquad \text{(S1)}$$

where $I_0$ is the integrated fluorescence intensity obtained at the critical angle for the TIR illumination, $\lambda$ is the wavelength of the illuminating light, and $n_1$ and $n_2$ denote the refractive indices of the solution and the glass coverslip, respectively. We fitted the TIR illumination angle-dependent integrated fluorescence intensity using the following equation:

$$I = I_0\left(\frac{\lambda}{4\pi\sqrt{n_1^2\sin^2\theta - n_2^2}}\right) + bg \qquad \text{(S2)}$$

where *bg* is the background signal on the measured fluorescence images.

### Supporting Figures

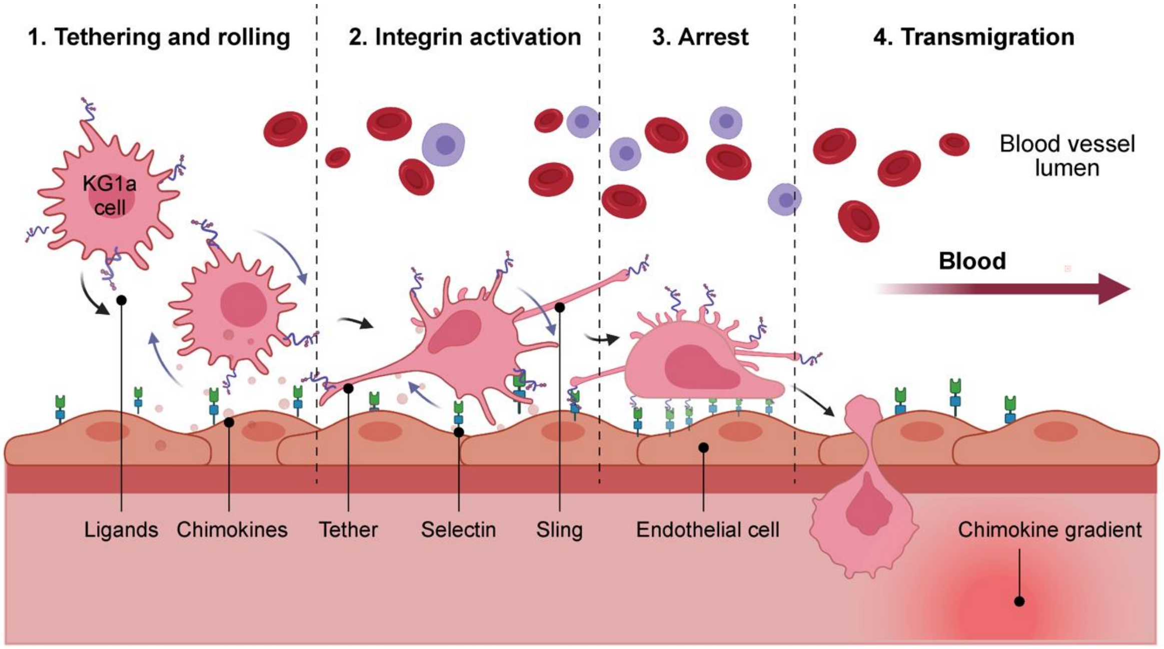


**Figure S1.** Schematic overview of the hematopoietic stem/progenitor cell (HSPC) homing mechanism. HSPC homing consists of five sequential steps: (1) Tethering and rolling, initiated by the binding of selectins expressed on endothelial cells to selectin ligands on the rolling HSPCs. (2) Activation, triggered by inflammatory signals, leading to cellular responses. (3) Arrest, facilitated by integrin binding, resulting in firm adhesion. (4) Transmigration, where HSPCs migrate through the endothelial layer and basement membrane. (5) Directed migration, where HSPCs move toward chemokine gradients to reach their target niche.

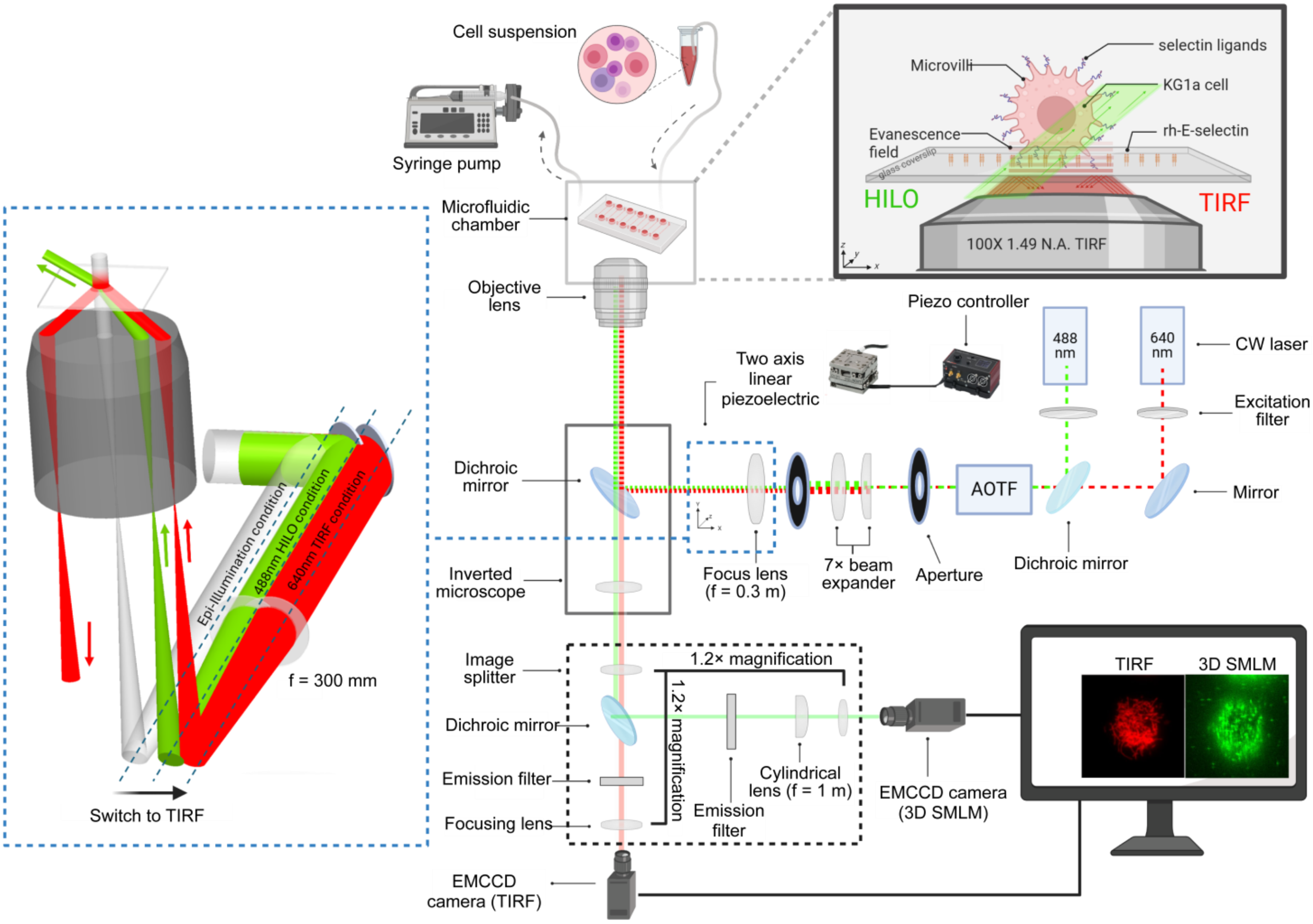


**Figure S2.** Schematic diagram of a combined TIRFM and 3D-SMLM setup. The setup allows reliable and reversible switching between total internal reflection (TIR) illumination and highly inclined and laminated optical sheet (HILO) illumination. Microfluidic devices required for the cell rolling assay are integrated into the microscope so that the combined TIRFM and 3D-SMLM imaging experiment can be conducted on the rolled cells.

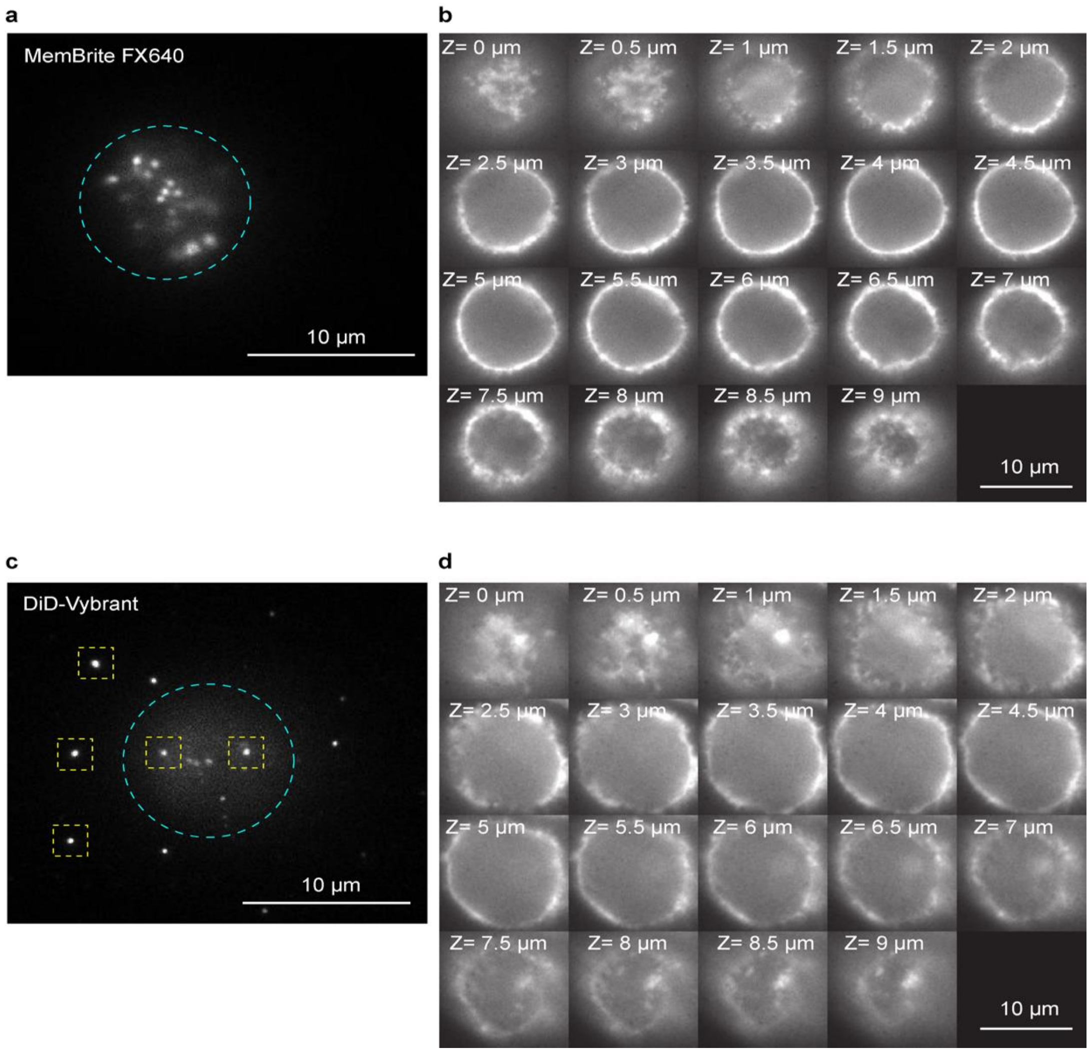


**Figure S3.** Uniform labeling of the cell membrane using membrane stains. (a) TIRF image of the bottom surface of a KG1a cell stained by MemBrite FX640 stain. The cyan circle indicates the cell surface region. (b) Fluorescence images of KG1a cells stained with MemBrite FX640 captured at varied Z-axis positions using wide-field epi-illumination fluorescence microscopy. (c) TIRF image of the bottom surface of a KG1a cell stained by DiD-Vybrant stain. The cyan circle indicates the cell surface region. Yellow squares show dye aggregates. (d) Fluorescence images of KG1a cells stained with DiD-Vybrant captured at varied Z-axis positions using wide-field epi-illumination fluorescence microscopy. The Z-axis values represent the distances from the coverslip surface.

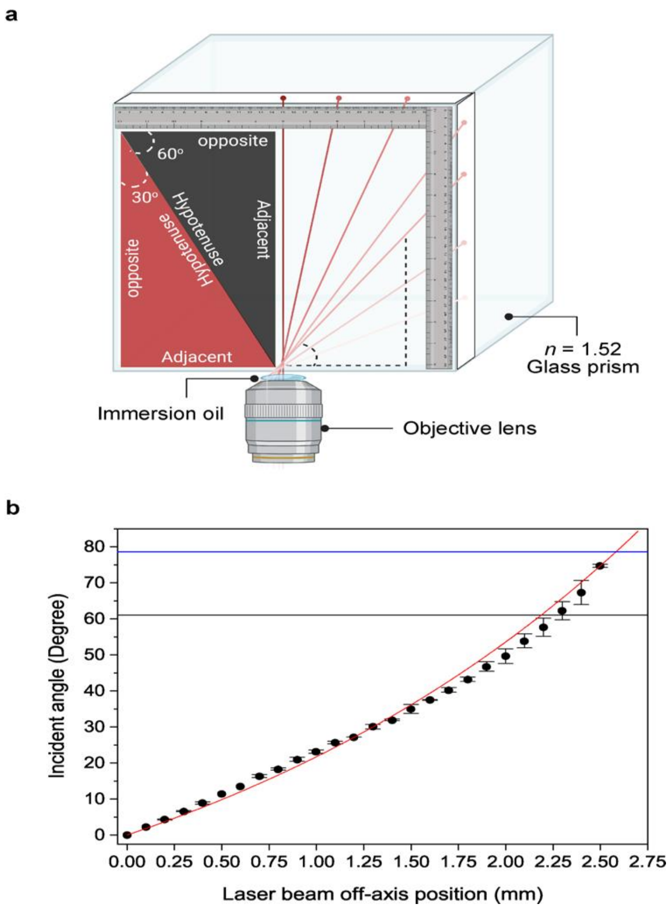


**Figure S4.** Determination of the incident angles. (a) Schematic diagram of the experimental setup for determining incident angles, where a glass prism with a refractive index of 1.52 is mounted on an inverted microscope stage in direct contact with a TIRF objective lens (N.A. = 1.49). An oil immersion layer was applied between the objective and the prism to match the refractive index. Incident angles were experimentally determined using trigonometric functions. (b) Experimentally determined incident angles at varied laser beam off-axis positions at the back focal plane of the objective lens. The black horizontal line represents the theoretically calculated critical angle for TIR illumination with the TIRF objective lens (61°, see Supporting Text 1). The blue horizontal line represents the maximum TIR angle achievable using the TIRF objective lens (79°, see Supplementary Text 1). The red line shows a single-exponential fitting.

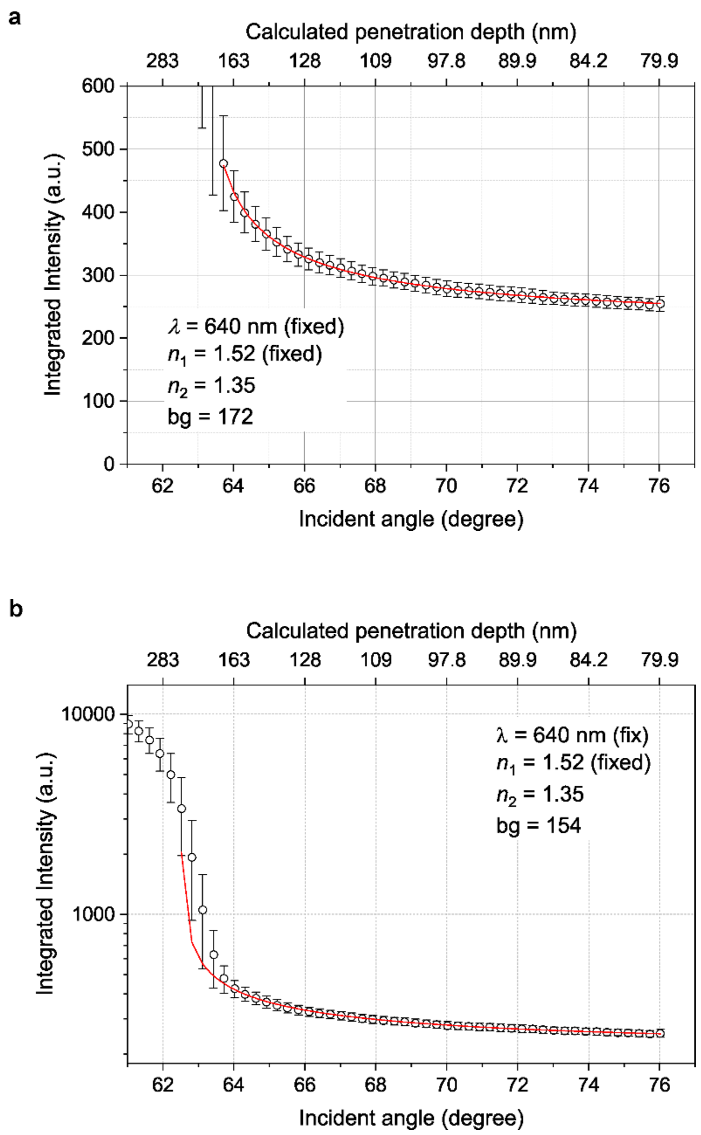


**Figure S5.** Verification of the incident angles and penetration depths of the excitation light. Mean integrated intensity profile of the 10 μg mL-1 AF640 fluorophores in an aqueous solution, measured across different TIR illumination angles. The error bars show standard deviations of 3 independent experiments. Red lines show fittings to equation S2 (see Supplementary Text 2) where λ is the wavelength of the excitation light, n1 and n2 denote the refractive index of the glass coverslip and the solution, and bg is the background signal in the imaging experiments. The range of the incident angle fitted to equation S2 was (a) 63.7° - 78° and (b) 62.4° - 78°.

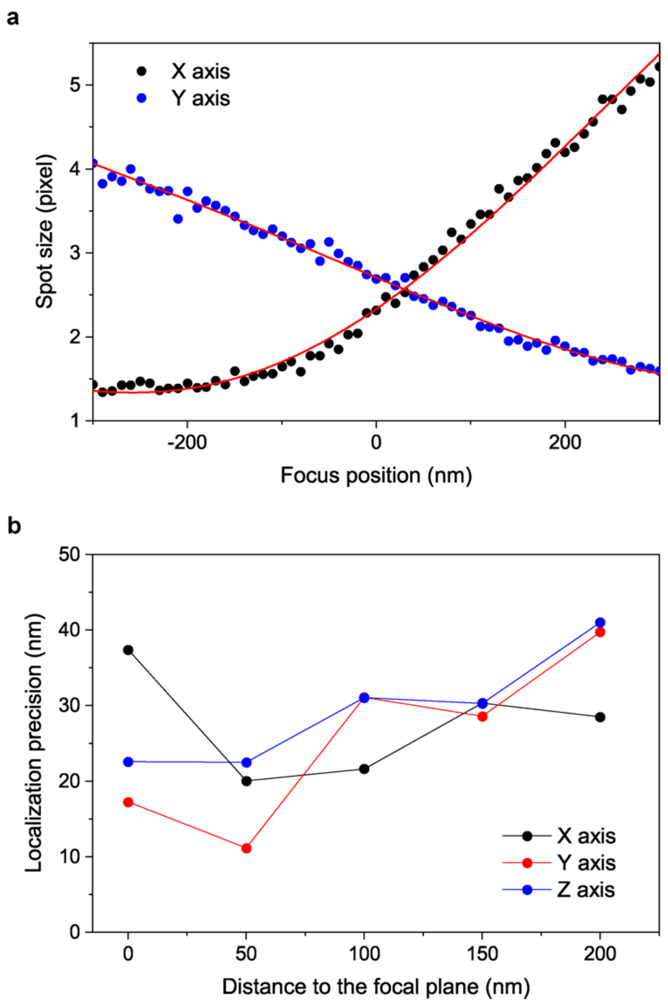


**Figure S6.** Localization precision of 3D-SMLM. (a) Z-position-dependent widths (standard deviations of the 2D Gaussian used for the fitting) of the fluorescent spots along the X- and Y-axes obtained for TetraSpeck fluorescent nanospheres (100 nm diameter). The red lines show fitting to polynomial functions, which were used for the Z-axis localization of the fluorescent molecules. (b) Localization precisions along X-, Y-, and Z-axes (standard deviations of the XYZ coordinates localized by 3D-SMLM) at varied distances from the focal plane. Localization data were obtained using 20 nm TetraSpeck fluorescent nanospheres captured over 10,000 frames.

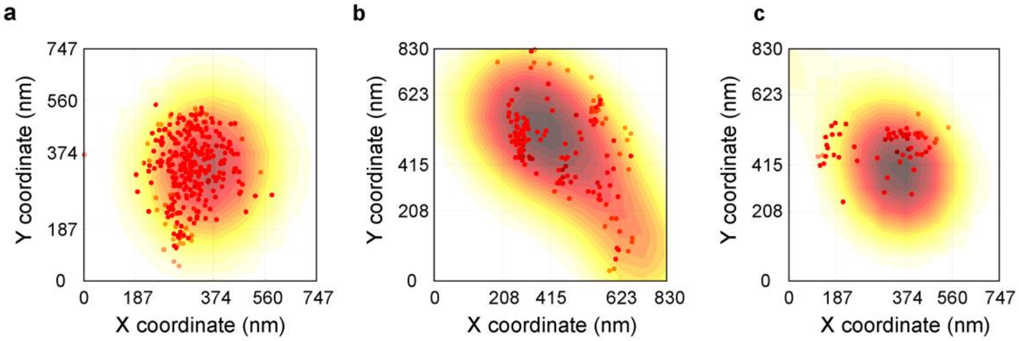


**Figure S7.** Spatial distribution of adhesion molecules on microvilli projected on the XY plane. XY projections of 3D single molecule localizations of (a) actin, (b) CD44, and (c) PSGL-1 on KG1a cells determined by 3D SMLM overlaid with a 3D cell surface topography image acquired by TIRF microscopy with MemBrite FX640 stain. The actin, CD44, and PSGL-1 were labeled by AF-488-conjugated phalloidin (actin) or immunolabeled with AF-488-conjugated antibodies (CD44 and PSGL-1). (a), (b), and (c) are the XY projections of the 3D images shown in Figure 5b, 6a, and 6e.

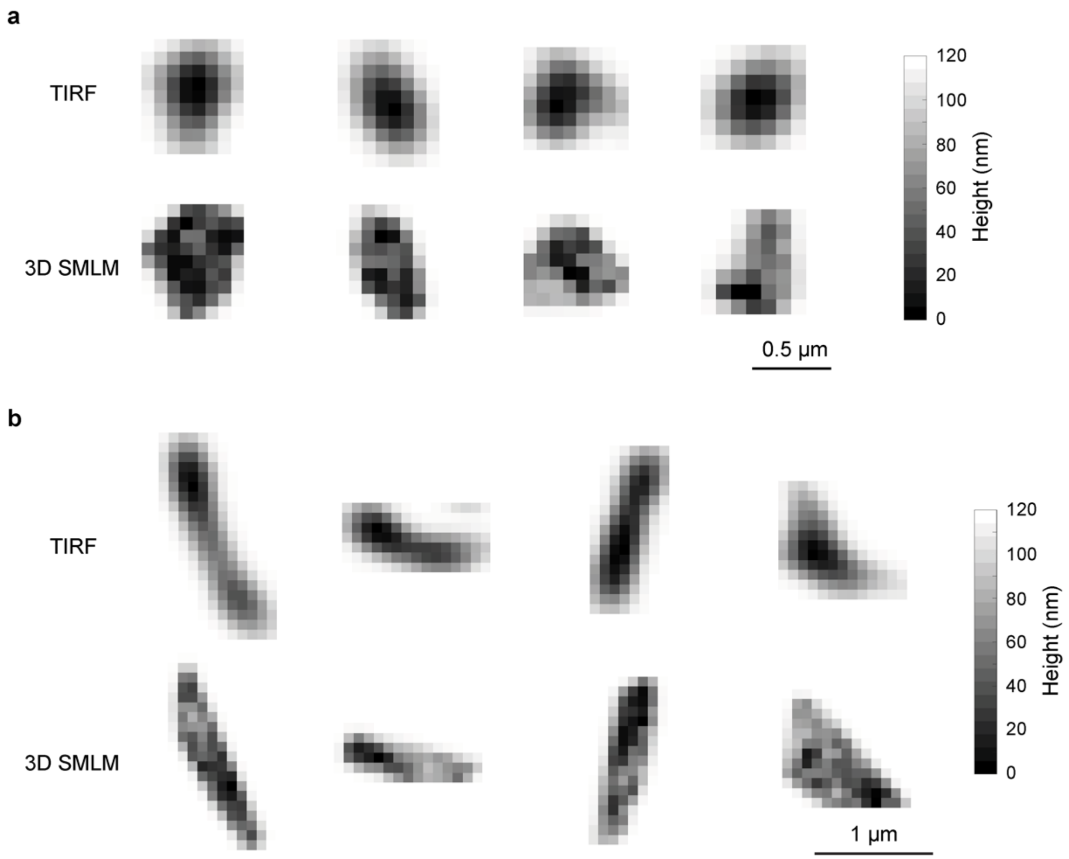


**Figure S8.** Comparison of the 3D topographic image of the cell surface and the 3D spatial boundary obtained from the 3D coordinates of individual actin molecules. Examples of normalized 8-bit 2D images obtained from the 3D topography of a single microvillus (top) and 3D spatial boundary obtained from the 3D coordinates of individual actin molecules (bottom). The axial distance from the coverslip surface is shown in grayscale. (a) Vertically oriented and (b) horizontally oriented microvilli were captured in the imaging experiment.

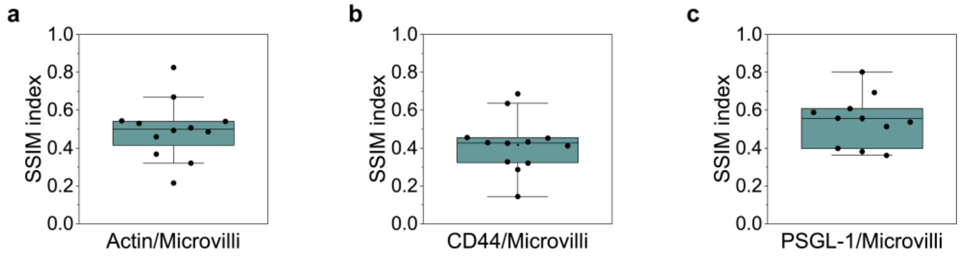


**Figure S9.** Structural similarity (SSIM) index analysis of the 3D topography of the microvilli generated by TIRFM and the 3D coordinates of the (a) actin, (b) CD44, and (c) PSGL-1 molecules determined by 3D-SMLM.

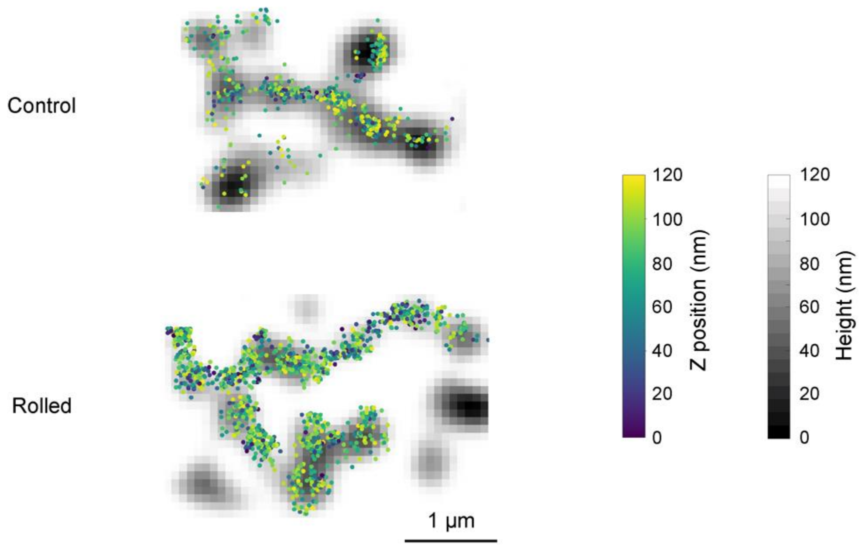


**Figure S10.** Effect of cell rolling on the architecture of microvilli and spatial distribution of actin. The grayscale cell surface topography images obtained by TIRFM are overlaid with the 3D coordinates of actin determined by 3D-SMLM. The grayscale shows the distance from the coverslip surface, and the color scale represents the Z-position of the actin molecules. The top and bottom panels show examples obtained for control and rolled cells over E-selectin.

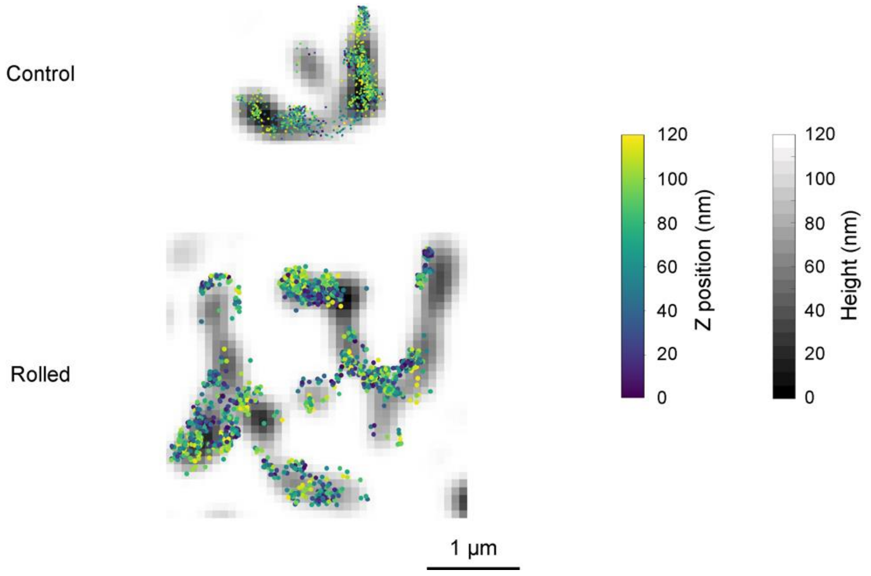


**Figure S11.** Effect of cell rolling on the architecture of microvilli and spatial distribution of CD44. The grayscale cell surface topography images obtained by TIRFM are overlaid with the 3D coordinates of CD44 determined by 3D-SMLM. The grayscale shows the distance from the coverslip surface, and the color scale represents the Z-position of the CD44 molecules. The top and bottom panels show examples obtained for control and rolled cells over E-selectin.

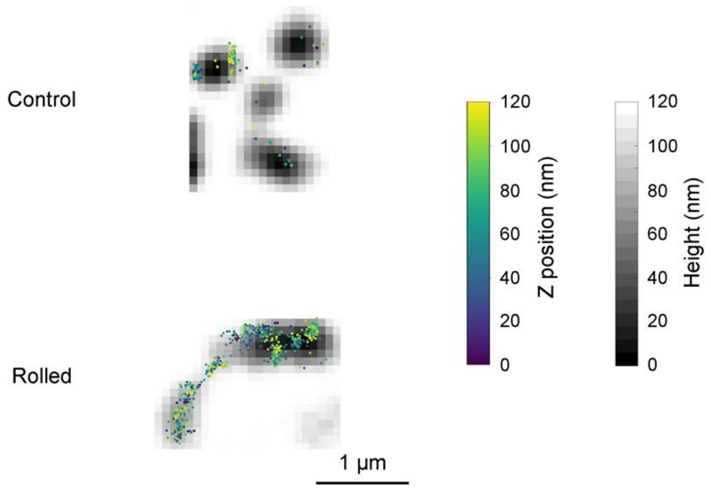


**Figure S12.** Effect of cell rolling on the architecture of microvilli and spatial distribution of PSGL-1. The grayscale cell surface topography images obtained by TIRFM are overlaid with the 3D coordinates of PSGL-1 determined by 3D-SMLM. The grayscale shows the distance from the coverslip surface, and the color scale represents the Z-position of the PSGL-1 molecules. The top and bottom panels show examples obtained for control and rolled cells over E-selectin.

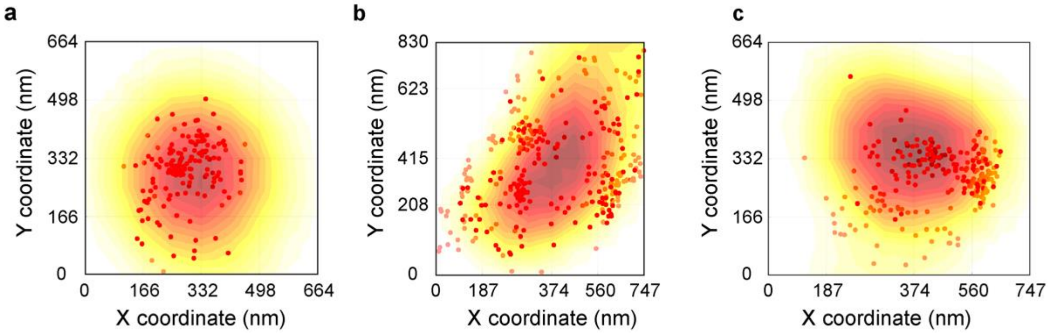


**Figure S13.** Spatial distribution of adhesion molecules on microvilli on the rolled cells projected on the XY plane. XY projections of 3D single molecule localizations of (a) actin, (b) CD44, and (c) PSGL-1 on KG1a cells rolled over E-selectin determined by 3D-SMLM overlaid with a 3D cell surface topography image acquired by TIRFM with MemBrite FX640 stain. The actin, CD44, and PSGL-1 were labeled by AF-488-conjugated phalloidin (actin) or immunolabeled with AF-488-conjugated antibodies (CD44 and PSGL-1). (a), (b) and (c) are the XY projections of the 3D images shown in Figure 7b, 7e, and 7h.
